# A comprehensive computational analysis investigating the relationships between phage codon usage, infection style, and number of tRNA genes

**DOI:** 10.64898/2026.03.19.712862

**Authors:** Nykki D. Ross, Sarah M. Doore

## Abstract

It has been known for decades that bacteriophages encode tRNA genes, but their function and the factors contributing to their acquisition and retention are unclear. Although tRNAs are found in a variety of phages infecting a variety of bacteria, many large-scale computational studies investigating tRNA acquisition and retention in phages are specific to *Mycobacterium* phages; however, these findings may not be representative of other phages or bacteria. This work uses a broader sampling of phages and hosts to investigate the relationships between codon usage bias, infection cycle, and tRNA gene numbers in phage genomes. We analyzed 154 phages infecting 7 host genera, including Gram-negative (*Escherichia*, *Shigella*, *Salmonella*) and Gram-positive (*Bacillus*, *Lactobacillus*, *Staphylococcus*, *Mycobacterium*) bacteria. Phages included temperate and virulent representatives, plus a range of tRNA numbers and morphologies. All phages and hosts were analyzed using four metrics: GC content, Effective Number of Codons, Relative Synonymous Codon Usage, and tRNA Adaptation Index. On a global scale, virulent phages with many tRNA genes show greater differences in codon usage and codon adaptation compared to their respective hosts. Gram-negative bacteria and their phages generally exhibit greater differences in codon usage compared to Gram-positive bacteria and their phages. Phages infecting Gram-negative hosts also tend to encode more tRNA genes. In nearly all genus-level comparisons, *Mycobacterium* phages were different from any other host and from global patterns. This suggests previous computational studies performed in *Mycobacterium* phages are likely not applicable on a global scale or to phages infecting other host genera.

**AUTHOR SUMMARY:** Bacteriophages, or phages, are viruses infecting bacteria. They are abundant in all environments, yet how they interact with their bacterial hosts is still not well-understood. Like other viruses, phages must rely on the host translational components to replicate and form new phage particles; and similarly to other parasites, phages have genomes that differ significantly from their hosts in terms of composition. In this work, we explore the relationship between phage lifestyle, number of tRNA genes encoded, and genome differences from the host using a variety of phages and their associated hosts. Phages can be either virulent (do not integrate into the host genome) or temperate (capable of integrating into the host genome), with differences from the host genome more pronounced in virulent phages. There are many phages that also carry tRNA genes, and having higher numbers of tRNAs is associated with larger differences from the host genome. The findings here indicate that virulent phages carrying large numbers of tRNAs diverge the most from host genome composition.

## BACKGROUND

Bacteriophages, or phages, are viruses infecting bacteria. They are present in high abundance in the environment, with phages found in every ecological niche in which bacteria are found at an approximate ratio of 10:1 phage particles to bacterial cells (1). Transfer RNAs (tRNAs) were first discovered in bacteriophage genomes in 1968 with the identification of 8 tRNA genes encoded together in a region of the *Escherichia* phage T4 genome (2). Since their first discovery, many bacteriophages have been shown to encode tRNA genes, though their functions are still poorly understood. Phages can encode anywhere from one to several dozen tRNA genes (3), though the reason for this variation is not known. There have been several hypotheses proposed explaining the presence of phage-encoded tRNAs, including the codon usage bias, host expansion, all-destructive phenotype, and antiphage defense escape hypotheses. Of these, the codon usage bias hypothesis is the most widely-accepted theory (4–8).

Codon usage bias refers to the unequal usage of synonymous codons for an amino acid, and is present across all domains of life (9–11). Codon biases are associated with gene expression levels and rates of translation, with highly-expressed genes showing higher levels of codon bias. Increased codon usage bias is also associated with increased translational efficiency, prioritizing codons that match tRNAs that are abundant in the tRNA pool of an organism (11,12). Because phages generally must rely on the host tRNA pool to translate their own proteins, one might assume that the phage codon usage would match the usage of its host. This is not always the case, though, and phages often exhibit codon biases more similar to those of other parasites with low GC content and AT-rich genomes (13). It is thought that phages encoding tRNAs do so in order to add to the host tRNA pool and supplement it with tRNAs that match codons used frequently by the phage but infrequently in the host. A second hypothesis stating expanded host ranges contribute to tRNA acquisition in phages arose from the codon usage bias hypothesis. In addition to increasing translational efficiency of phage proteins, the tRNAs encoded by a phage can also expand its host range by allowing the phage to infect hosts with varying codon biases and different GC content (5,14).

In some cases, while codon usage bias differences may be present between the phage and the host throughout the genome, these differences are more pronounced in the late genes of the phage, which encode the virion structural components. In the all-destructive infection hypothesis, it is stated that phage-encoded tRNAs are needed specifically in the late stages of infection to translate the structural genes following the degradation of the host genome early on in infection (15,16). When the host genome is degraded, the tRNA genes encoding host tRNAs are also degraded, halting the transcription of the tRNAs. Once the host tRNA pool is depleted, the phage must rely on its own tRNAs to continue translation.

It was also discovered that in some *Mycobacterium* phages, the anticodon loops of phage-encoded tRNAs contain mutations that render them insensitive to host anticodon nucleases (17). Anticodon nucleases are a defense mechanism employed by bacteria upon phage infection, degrading the host tRNAs in order to reduce the resources available to the phage for replication (18). In the case of the nuclease escape hypothesis, the anticodon loop mutations in the phage-encoded tRNAs allow them to evade this host defense mechanism, permitting translation of phage proteins even as host tRNAs are depleted.

More recently, phage-encoded tRNAs have also been shown to assist in the evasion of other antiphage defense mechanisms. The phage anti-restriction induced system (PARIS) targets the anticodon stem-loop of tRNA-Lys-TTT in order to prevent coinfection of a host cell with other phages (19,20). The *Escherichia* phage T5 encodes a highly-expressed tRNA-Lys-TTT to counteract the effects of PARIS and allow T5 infection to proceed even in the presence of other phages. T5 tRNAs have also been implicated in the evasion of host retrons (21), namely the *E. coli* retron Eco7, which degrades the host tRNA-Tyr-GTA via two effector proteins, PtuA and PtuB (22). T5 encodes a highly-expressed tRNA-Tyr-GTA, which is thought to compensate for this degradation.

Additionally, the purpose of phage-encoded tRNAs can depend on lifestyle. A phage can have one of two lifestyles. Virulent phages do not integrate into the host genome and are only capable of lytic infection, where the phage replicates inside the host cell and bursts the cell to release new phage particles to spread to other host cells. Temperate phages are capable of lysogeny, in which the phage either exists as an extrachromosomal element or integrates into the host genome and exists as a prophage. Some temperate phages encode host virulence factors in their genomes (23), while others encode serotype conversion genes that are capable of altering the serotype of the host (24–27). The lytic cycle of prophages can be induced by environmental stressors, specifically by DNA damage like that triggered by mitomycin C treatment (28). In temperate phages, tRNAs have been shown to serve as integration sites during lysogeny (29). In the case of tRNA-dependent lysogeny, the integration site of a phage interrupts a host tRNA gene, but the common core of the phage and host attachment sites (attP and attB, respectively) is too short to reconstruct the host tRNA. Instead, the temperate phage encodes a tRNA of the same isotype in its genome, compensating for the loss of the host tRNA.

Whatever the mechanisms and effects, the lifestyle of a phage and its codon usage bias seem to have a role in phage tRNA acquisition and retention, though this complex relationship is not well-understood. This work investigates these relationships by examining the effects of lifestyle and number of phage-encoded tRNAs on codon usage bias and how the codon usage bias of phages differs from that of their hosts. A total of 154 phages of diverse taxonomy infecting 7 host genera were used in this comprehensive analysis, including both temperate and virulent phages, as well as phages encoding anywhere from zero to 37 tRNA genes. The host genera in this study include both Gram-negative (*Escherichia*, *Shigella*, and *Salmonella*) and Gram-positive (*Bacillus*, *Lactobacillus*, *Staphylococcus*, and *Mycobacterium*) bacteria to assess both global impacts of lifestyle and tRNA genes on codon usage, as well as host-specific impacts.

The majority of the large-scale studies on tRNAs in phages have been performed using Mycobacterium phages due to the large number of these phages which have been isolated and characterized (4,17,30). However, the results of this study indicate that the data gathered from researching *Mycobacterium* phages may not be an accurate representation of trends in other organisms, nor an accurate representation of global trends. When investigating the GC content, relative synonymous codon usage (RSCU), and tRNA adaptation index (tAI) of all phages compared to their hosts, the *Mycobacterium* phages consistently showed different patterns from the other host genera. Additionally, while global trends indicate that virulent phages with high numbers of tRNAs have the largest differences in codon bias compared to their hosts, variation is seen when looking at each host genus individually. In general though, phages infecting Gram-negative hosts tend to have larger differences in codon bias and higher numbers of tRNA genes. This indicates that the codon bias of these phages drives their tRNA acquisition, supporting the codon usage bias hypothesis. The results seen for phages infecting Gram-positive hosts show more variation and less support for codon bias driving the acquisition of their tRNAs. This study aims to fill this gap in knowledge by investigating diverse phages infecting a range of host bacteria both on the genus level and global scale.

## MATERIALS AND METHODS

### Selection of Phage Genomes

The National Center for Biotechnology Information (NCBI) and the PhageScope (31) database were used to identify virulent and temperate phages for all hosts. Phages must have had annotated genomes and must have the full genome sequences available in GenBank or RefSeq. Additionally, an emphasis was placed on phages with tRNA genes encoded in their genomes. A total of 154 phages were identified for this study, infecting 7 host genera: *Escherichia*, *Shigella*, *Salmonella*, *Bacillus*, *Staphylococcus*, *Lactobacillus*, and *Mycobacterium*. For most hosts, a single host strain was used, but in the case of *Bacillus* and *Lactobacillus*, it was not possible to find enough phages meeting the criteria. In these cases, more than one host strain was used. Additionally, as *Salmonella enterica* has many serovars, and no single serovar was infected by enough phages meeting our criteria, the two most common serovars (Typhimurium and Enteritidis) were included. For each host genus, a total of 17-24 phages were chosen.

### Host and Phage Genomes Used in the Comprehensive Analysis

The full genomes and coding sequences for the host species *Escherichia coli* K12 (NC_000913.3), *Shigella flexneri* 2457T (NC_004741.1), *Salmonella enterica* serovar Typhimurium SL1334 (NC_016810.1), *Salmonella enterica* serovar Enteritidis 92-0392 (NZ_CP018657.1), *Mycobacterium smegmatis* mc(2)155 (NC_008596.1), *Bacillus subtilis* 168 (NC_000964.3), *Bacillus cereus* ATCC 14579 (NZ_CP138336.1), *Staphylococcus aureus* NCTC 8325 (NC_007795.1), *Lactobacillus paracasei* 8700:2 (NC_022112.1), *Lactobacillus brevis* NCTC13768 (NZ_LS483405.1), and *Lactobacillus plantarum* WCFS1 (NC_004567.1) were obtained from the NCBI GenBank databases using their respective accession numbers. All tRNA genes were identified using the tRNAscan-SE 2.0 web program (32) in order to determine COVE scores for tRNA adaptation index (tAI) analysis.

The full genomes and coding sequences for all phages used in this analysis were also obtained from the NCBI GenBank database, and all phages with their respective accession numbers are listed in **Table 1**. When available, the RefSeq accession number is used; otherwise, the GenBank accession number is used. Phage tRNA genes were identified using the tRNAscan-SE 2.0 (32) web program and the ARAGORN (33) web program, as for some phages the programs gave slightly different results.

**Table 1.**
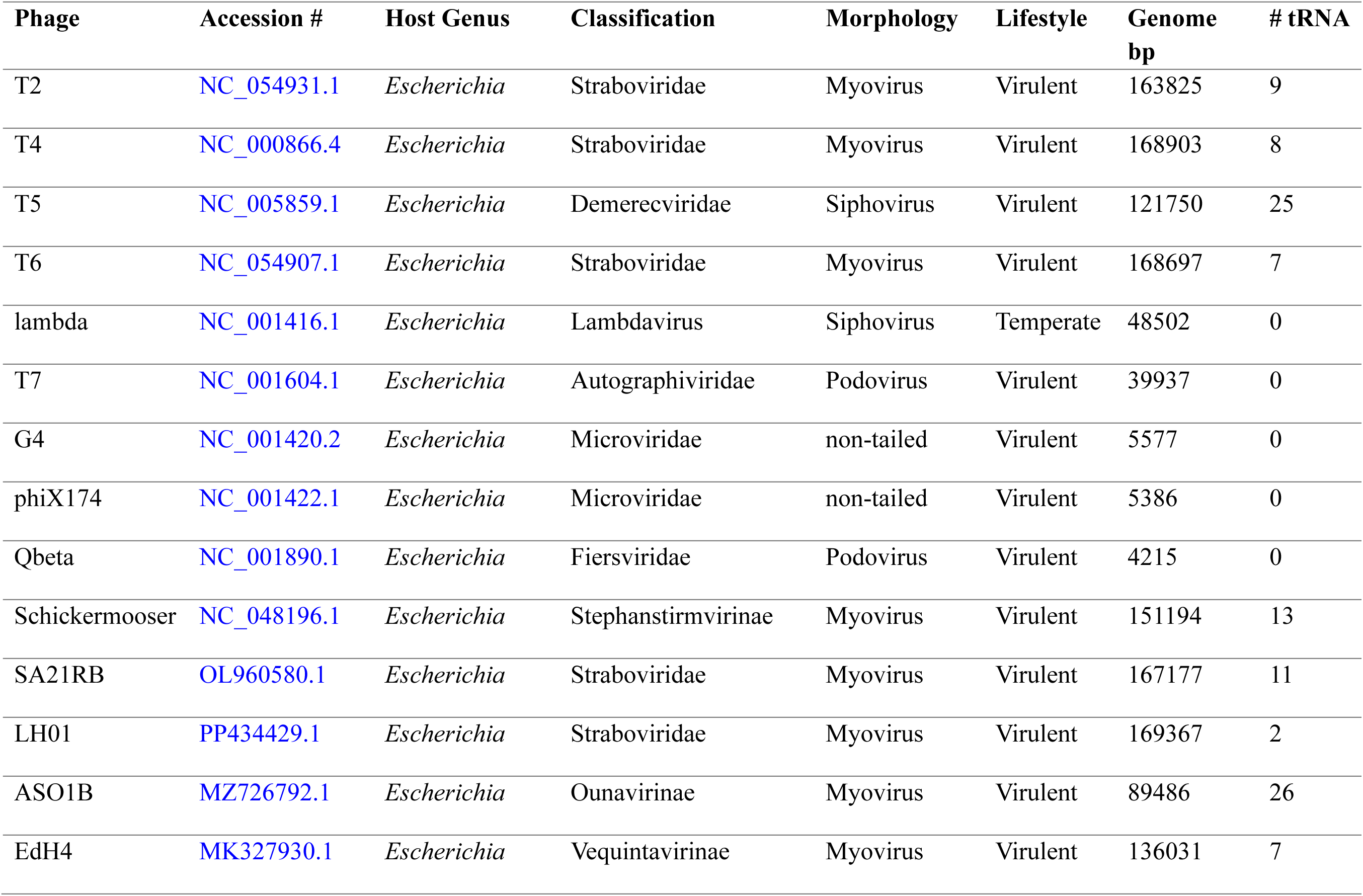
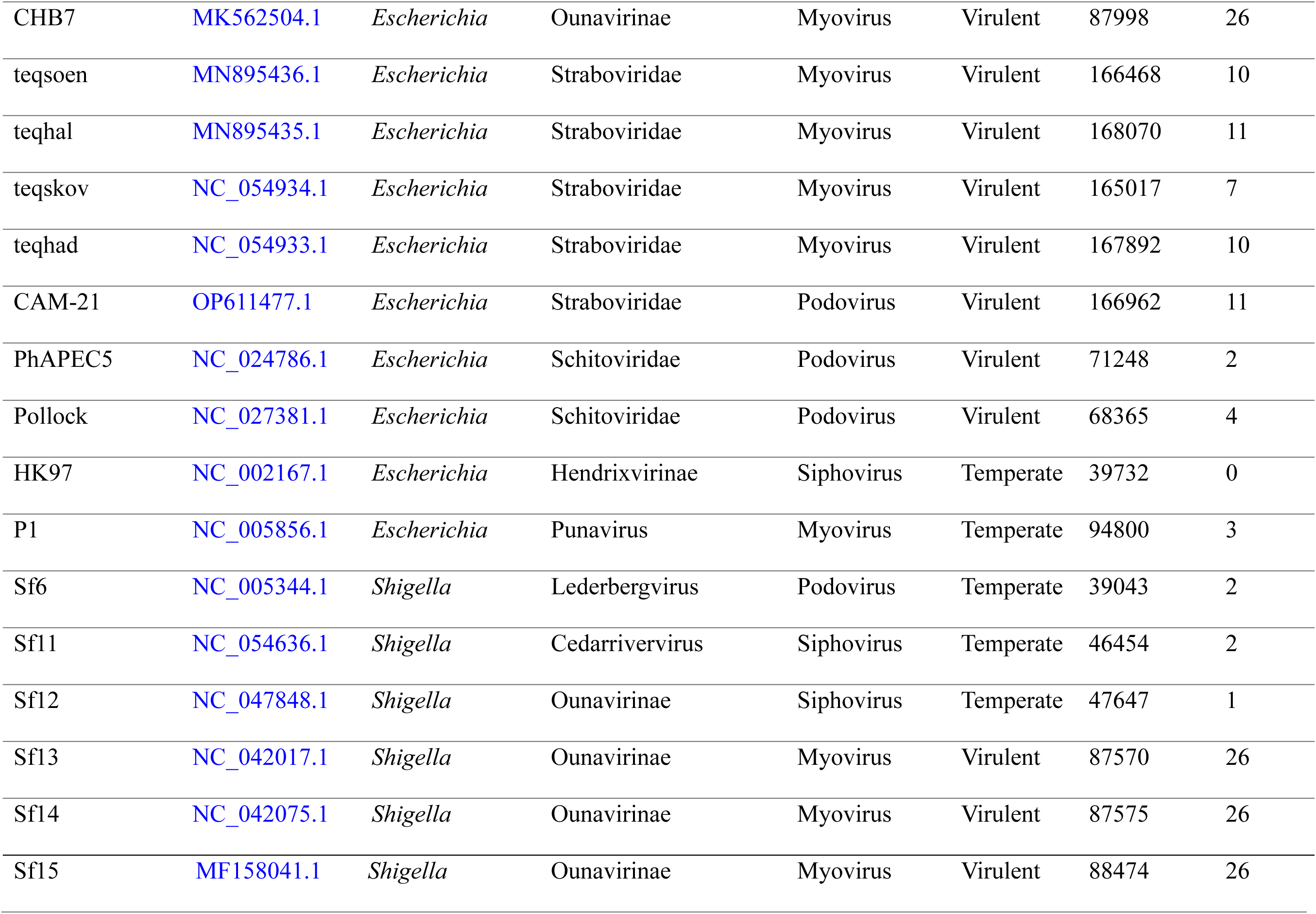
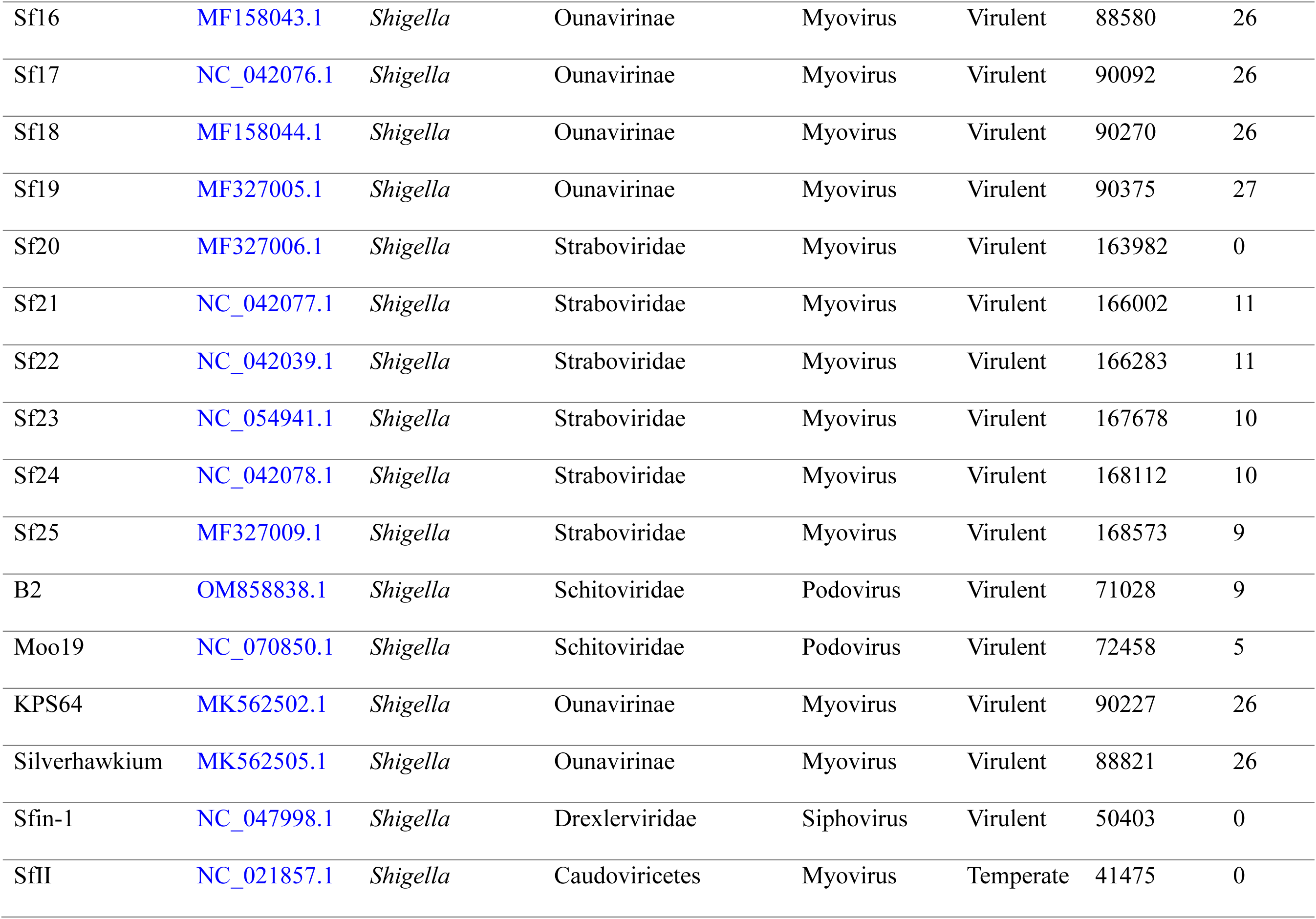
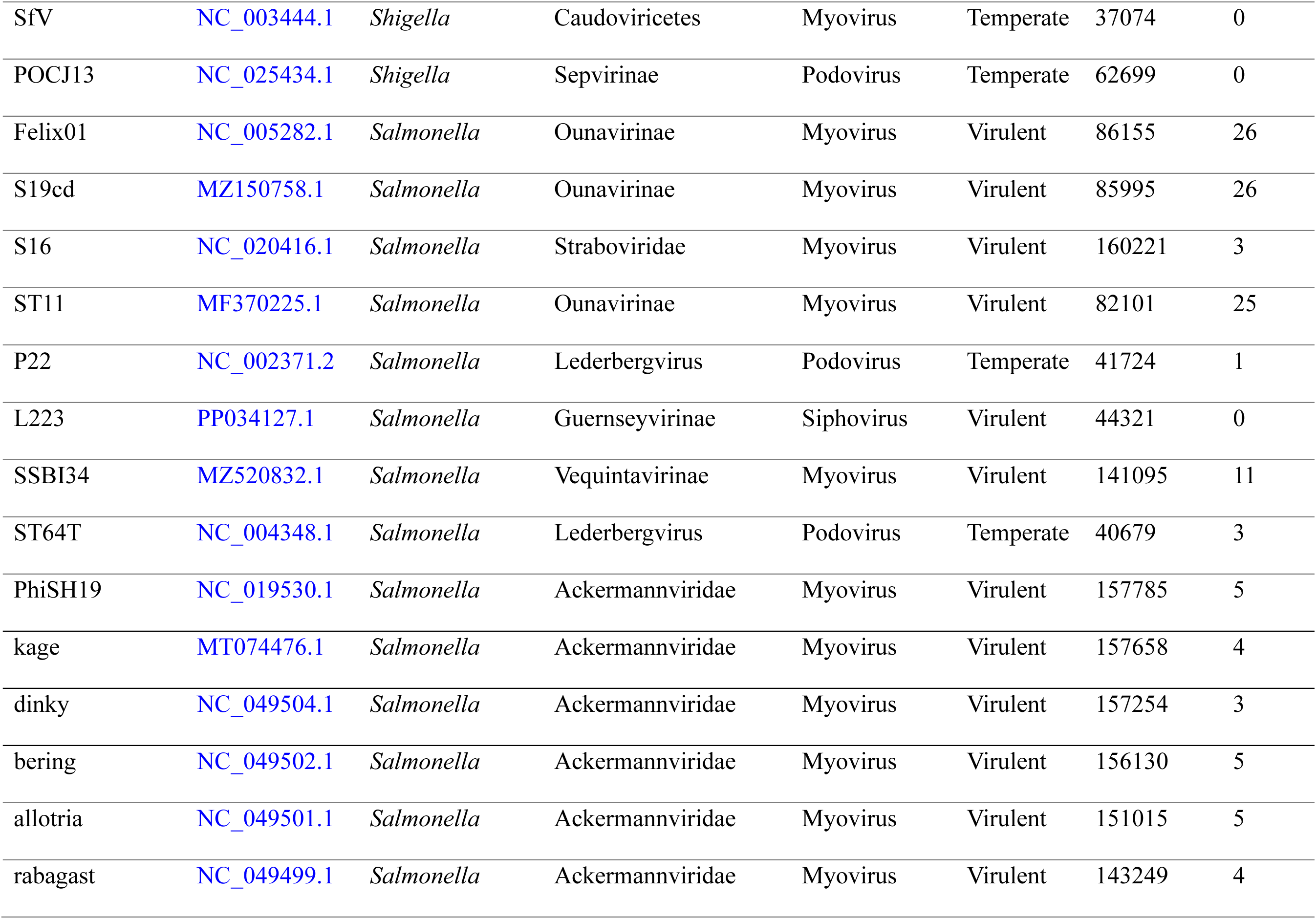
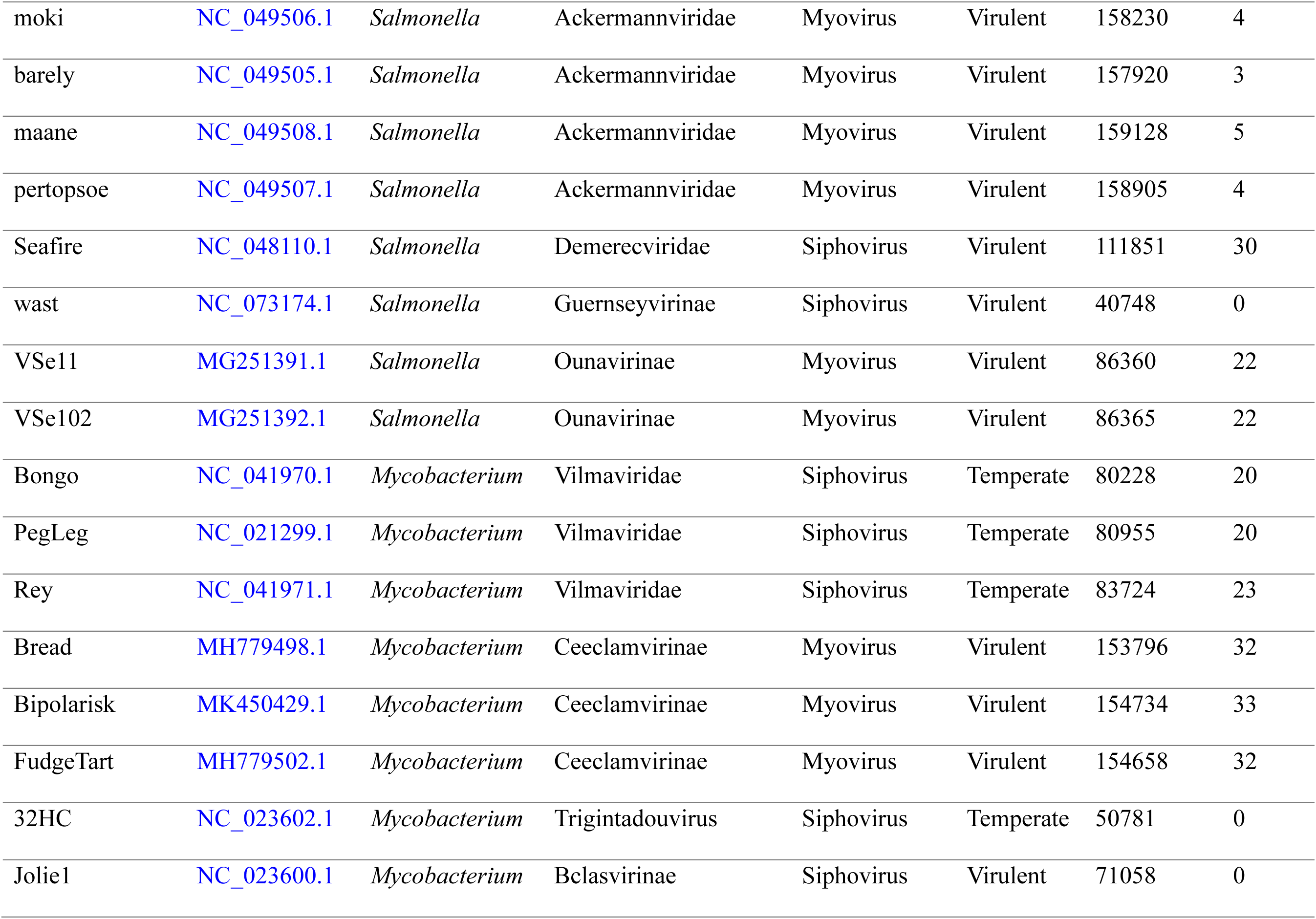
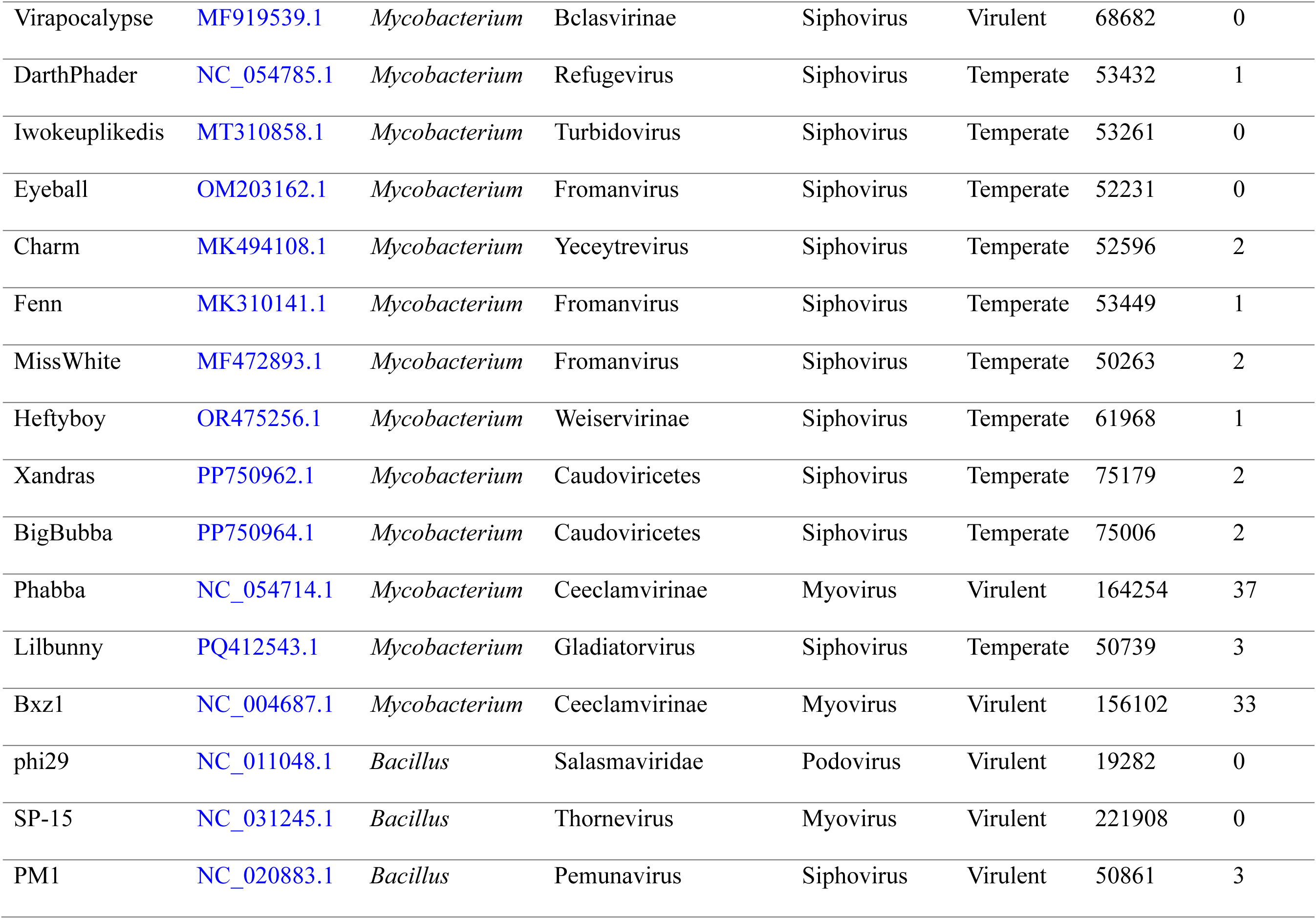
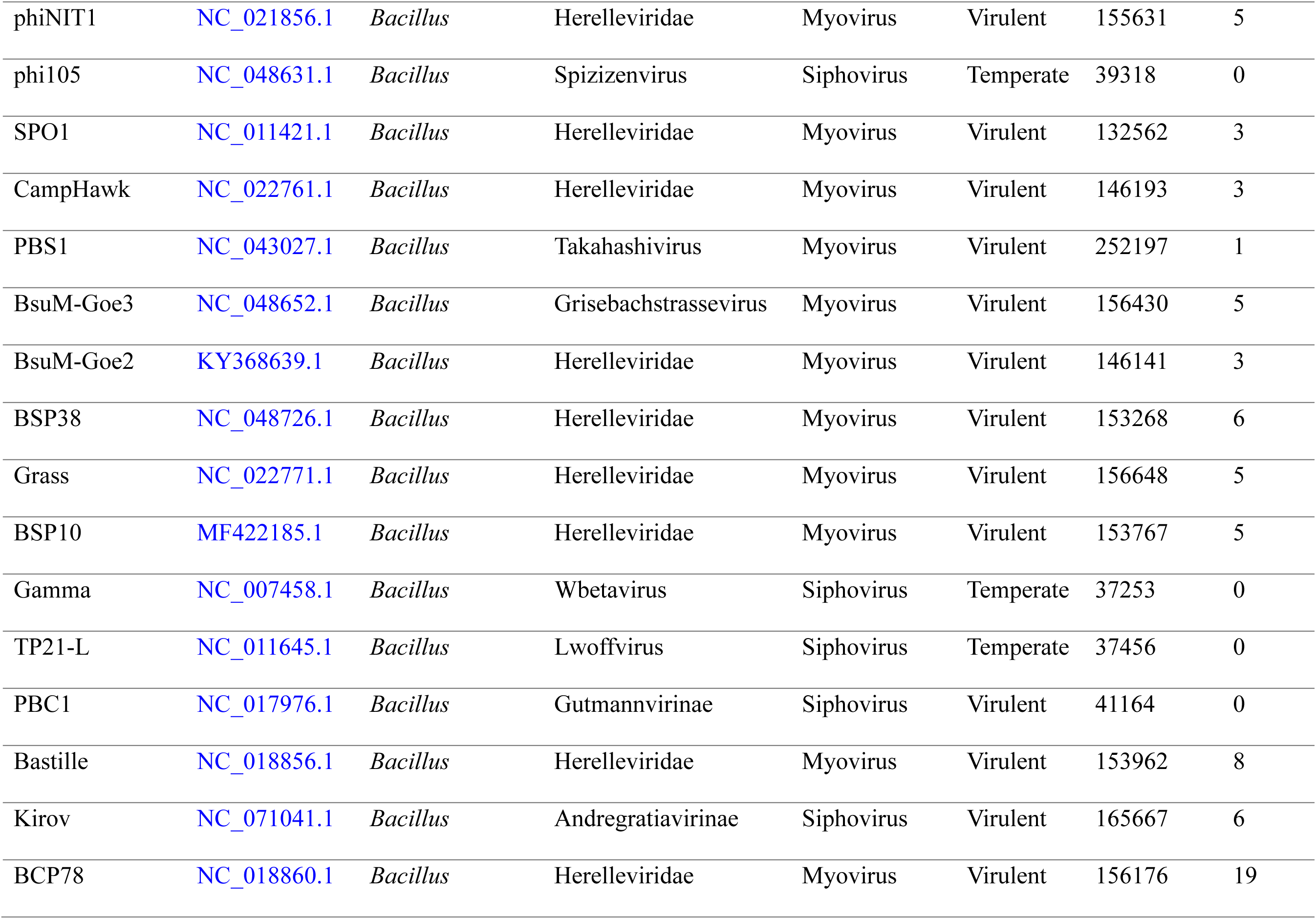
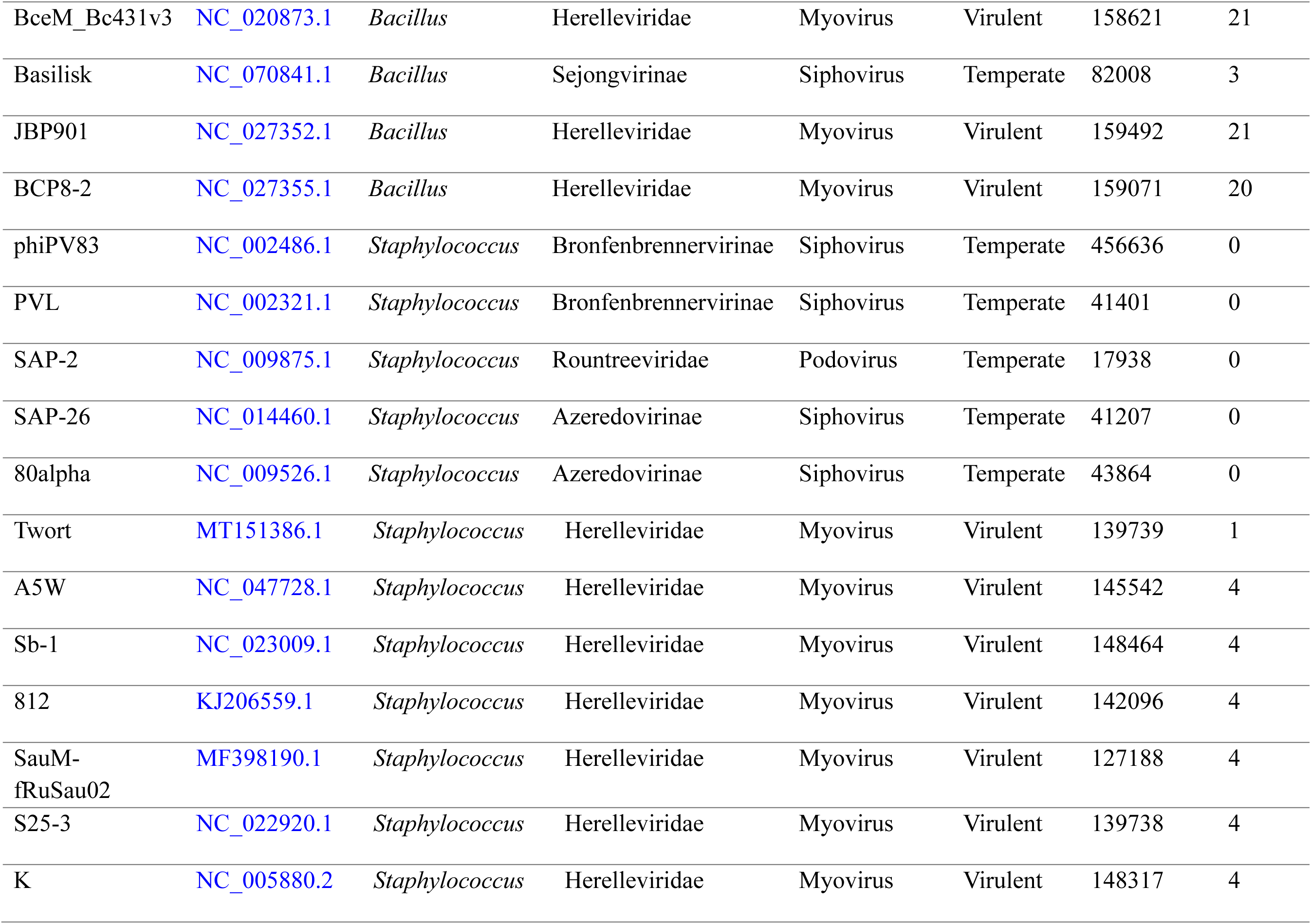
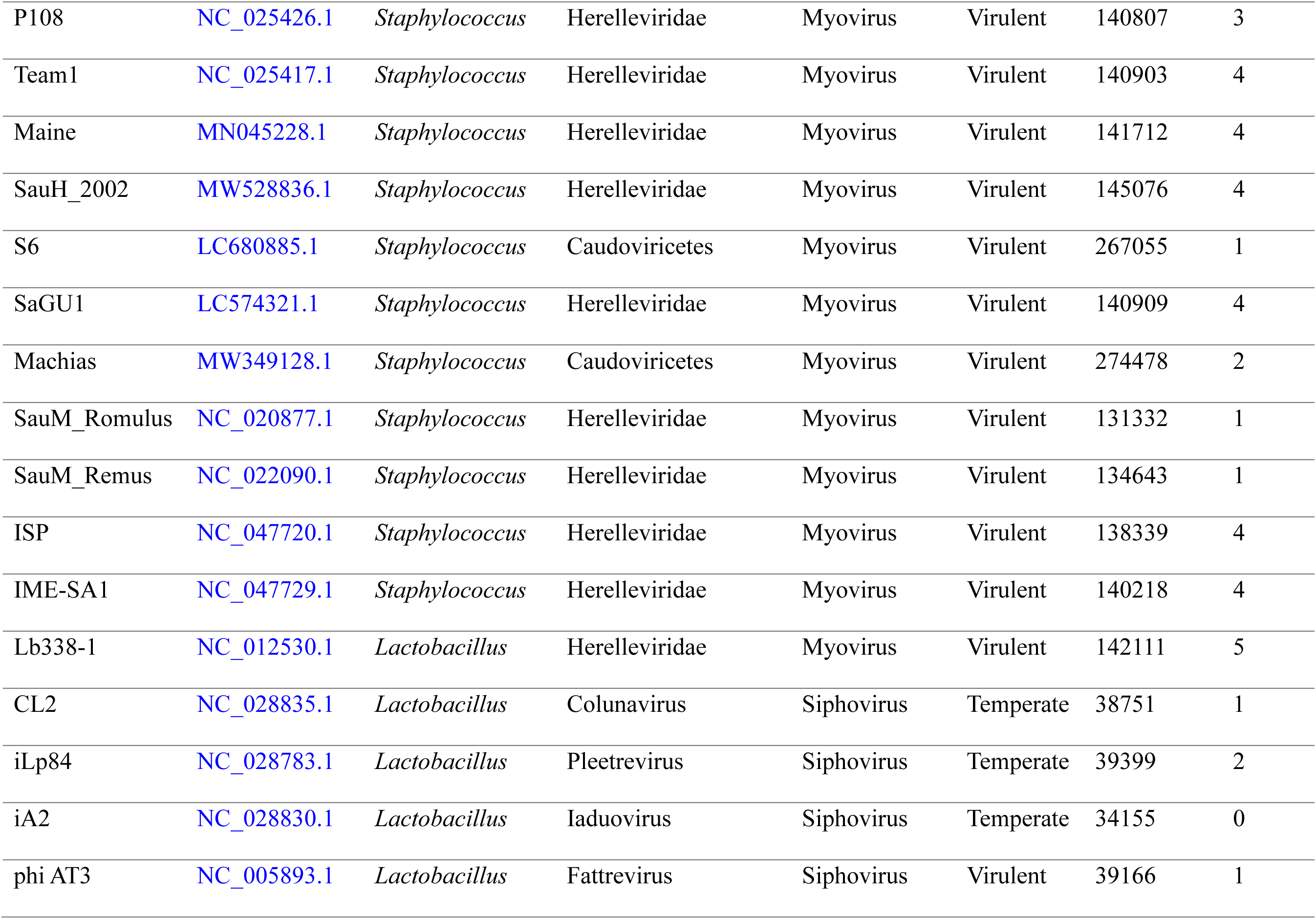
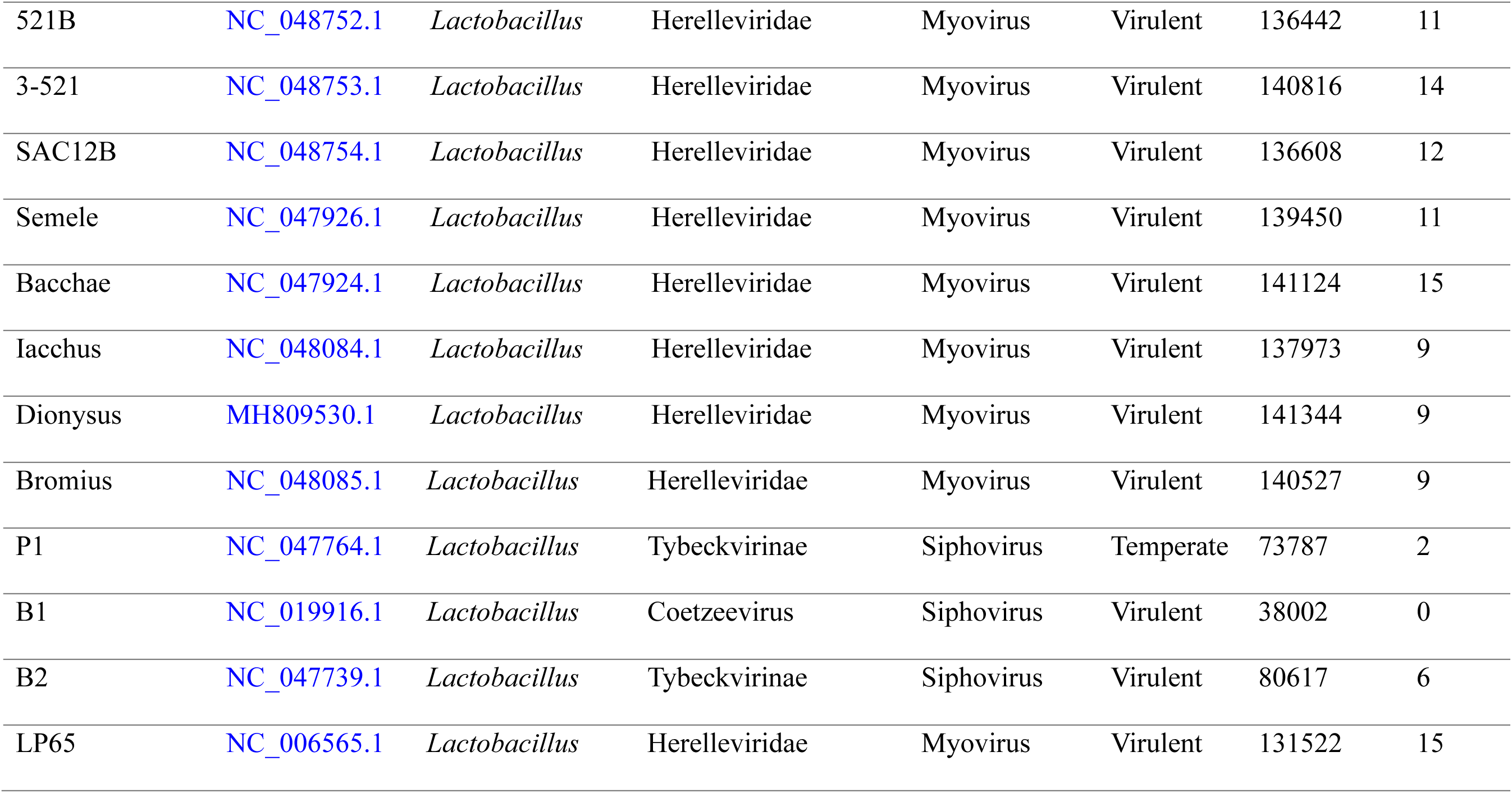
Characteristics of the phages used in analysis.

### Calculation of GC Content and ΔGC Content for Phages and their Hosts

The total GC content, as well as the codon position-specific GC content (GC1, GC2, and GC3, corresponding to codon positions 1, 2, and 3, respectively) were calculated and visualized in R using the readxl (34), dplyr (35), tidyr (36), purrr (37), stringr (38), seqinr (39), scales (40), and ggplot2 (41) packages. The GC content was calculated for all genes for all hosts and phages and visualized with violin plots. The mean GC content of each phage was compared to that of its respective host to determine the significance of the differences in genome composition. The ΔGC values were also calculated for total GC, as well as the codon position-specific GC content, by subtracting the mean host GC content for total GC, GC1, GC2, and GC3 from the mean GC content of the phage. The ΔGC content was then used for further comparisons between groups of phages within and between host genera, as this standardized the data to prevent biases that may occur with hosts exhibiting different GC content. Phage ΔGC values were compared based on lifestyle (Temperate vs. Virulent) and number of tRNA genes (High: > 5, Low: 1 – 5, None: 0) both within the host genus and across host genera, with the comparisons visualized with heatmaps.

### Calculation of the Effective Number of Codons (EnC) for Phages and their Hosts

The effective number of codons reflects the number of unique codons in a coding sequence, returning a single value for each gene. Values range from 20 to 61, with 20 meaning one codon is used per amino acid (highly biased) and 61 meaning all codons are used (low bias) (42). Lower EnC indicates possible selection mechanisms affecting codon usage that modify translational efficiency. Higher EnC values suggest the codon selection is due to genetic drift, rather than selection. EnC values were calculated per gene for all phages and their hosts in R using the packages readxl (34), dplyr (35), tidyr (36), purrr (37), readr (43), stringr (38), and seqinr (39), and the Nc plots were generated using the ggplot2 (41) package. EnC values for each phage were calculated and compared to the EnC values of their hosts. The EnC distance was then calculated (ΔEnC), which measures how different a phage’s EnC values are from those of its host. The ΔEnC was used to compare the phage tRNA groups (High, Low, and None) both within a host genus and across host genera, as well as comparing the lifestyle of phages (Temperate vs. Virulent) within a host genus and across host genera.

When compared to the GC3 content of an organism, the impact of selection and genetic drift on the codon usage of an organism can be determined using the Nc plot, which plots EnC against GC3 values (44). The EnC distance, which is the vertical distance of the actual gene EnC value from Wright’s curve (45), is calculated using the formula

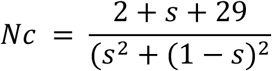

where *s* is the GC3 value. This generates a curve for the Nc plot. Values that fall below the curve indicate bias is likely due to selection pressures on expression levels, while values that fall on or above the expected curve indicate genetic drift determines the codon usage.

### Calculation of Relative Synonymous Codon Usage (RSCU) and ΔRSCU for Phages and their Hosts

The relative synonymous codon usage provides a measurement of how often specific synonymous codons are used for a given amino acid, calculated by determining the frequency of each synonymous codon in a genome or coding sequence (46). Values range from 0 (no usage) to the total number of synonymous codons for that amino acid. RSCU is calculated using the formula

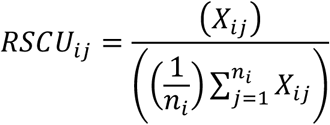

where *X*_{*ij*}_ is the frequency of the codon *j* for the *i^th^* amino acid. The number of synonymous codons for the *i^th^* amino acid is represented by *n_i_*. Calculations were performed in R using the packages dplyr (35), readr (43), tidyr (36), purrr (37), stringr (38), and forcats (47), and visualized using the packages ggplot2 (41) and pheatmap (48). RSCU values per codon were calculated for each coding sequence in each phage as well as each coding sequence in each host. The mean RSCU values for each codon in each phage were compared to the mean RSCU values of its respective host. ΔRSCU values were calculated by subtracting the host RSCU from the phage RSCU, and these values were subsequently used to compare codon usage between phage lifestyle groups and tRNA groups both within the host genus and across host genera. RSCU values were visualized using heatmaps.

### Calculation of tRNA Adaptation Index (tAI) and ΔtAI for Phages and their Hosts

The tRNA adaptation index estimates the translational efficiency of a gene based on how well the codon usage of that gene correlates with the tRNA pool of that organism. The tRNA gene sequences were extracted from each host and run through tRNAscan-SE 2.0 (32) to obtain the COVE scores, a measure of tRNA gene quality, for their respective tRNA pools. These scores were then used in the anticodon library generation during tAI analysis. The tAI values were calculated in R for all phages and their respective hosts with the host tRNA pools using the packages dplyr (35), readr (43), tidyr (36), purrr (37), stringr (38), and forcats (47) and visualized using the ggplot2 (41) package. Codon weights for tAI are calculated by matching codons to an organism-specific tRNA pool using the formula

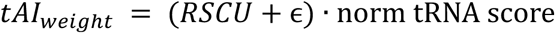

where ɛ is a small value (1.00 x 10^-06^) to prevent division by zero during log calculations (49), RSCU is the relative synonymous codon usage value for a given codon, and the norm tRNA score is the normalized score for the tRNA carrying the complementary anticodon. The geometric mean of all tAI weights is calculated using the formula

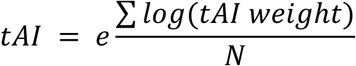

where N is the number of matched codons for a given anticodon. Finally, normalization of tAI values across a genome (so that the maximum value is 1) are calculated with the formula

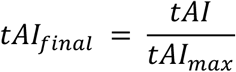

where tAI is the calculated tRNA adaptation index value for a given gene and max(tAI) is the maximum tAI across all genes in the genome.

The tAI values of phage genes were compared to those of their respective hosts, and the ΔtAI was calculated for each phage gene from these data by subtracting the mean host tAI values from the mean phage tAI values. ΔtAI was subsequently used to compare phages based on lifestyle and tRNA group both within the host genus and across host genera.

### Statistical Analyses

All statistical analyses were conducted in R using the packages glmmTMB (50), emmeans (51), rstatix (52), vegan (53), coin (54), FSA (55), effsize (56), broom (57), and broom.mixed (58) and visualized with the ggplot2 (41) package. To analyze the statistical significance of the difference in GC content between a phage and its respective host, the Wilcoxon rank-sum test was used to compare all host genes to all phage genes for total GC, GC1, GC2, and GC3. All ΔGC analyses include the Δtotal GC, ΔGC1, ΔGC2, and ΔGC3. To compare phage ΔGC content based on lifestyle, the Wilcoxon rank-sum test was also used. These comparisons included comparing phage lifestyles for each individual host, as well as comparing the ΔGC content based on phage lifestyle across all hosts. To compare the ΔGC content of the phage tRNA groups, the Kruskal-Wallis test was run for each host to compare phages with high numbers of tRNAs, low numbers of tRNAs, or no tRNAs, followed by Dunn’s post-hoc test with Bonferroni correction. Following the Kruskal-Wallis test, a pairwise Wilcoxon rank-sum test was run with Bonferroni correction to compare the tRNA groups High vs. Low, High vs. None, and Low vs. None. Lastly, a Wilcoxon rank-sum test was run to compare phages with tRNAs (High + Low) with phages without tRNAs (None). To compare the ΔGC content between phage tRNA groups across all genera, the same protocol as above was followed, but with data from all hosts pooled. Lastly, a linear mixed-effects model was used to test the effect of phage lifestyle across all hosts while controlling for host-to-host differences. This model used lifestyle as the fixed effect, and host as the random effect, following the model ΔGC ∼ lifestyle + (1|Host).

The Wilcoxon rank-sum test was used to compare the EnC of phages to the EnC of their hosts at the gene level. Likewise, for lifestyle comparisons, the Wilcoxon rank-sum test was also used to compare the ΔEnC of virulent and temperate phages both within each genus and globally. Effect size was calculated with Cliff’s delta test. For the tRNA group comparisons, the Kruskal-Wallis test was used to determine if there were significant differences in ΔEnC between phages with high numbers of tRNAs, those with low numbers of tRNAs, and those with no tRNAs. If the Kruskal-Wallis test showed significant results, Dunn’s post-hoc test with Benjamini-Hochberg (BH) correction for false discovery rate (FDR) was used to perform a pairwise comparison of the different tRNA groups. If only two tRNA groups existed in a genus, the Wilcoxon rank-sum test was used instead. Effect size was calculated using Cliff’s delta test. This was performed both at the host genus level and globally across all phages. The correlation between EnC and GC3 was calculated using Spearman’s correlation to determine where phage genes fall on Wright’s expected Nc curve.

The One-sample Wilcoxon signed rank test against a null hypothesis of µ = 0 was used to compare the RSCU values of each phage with those of its host on a per-codon basis. ΔRSCU analyses were also run on a per-codon basis to determine lifestyle-specific and tRNA group-specific codon usage patterns. A permutation-based Wilcoxon test with 10,000 resamples was used to compare the ΔRSCU per codon of phages based on lifestyle within each host genus, with Cliff’s delta test run to determine effect size, using a BH correction for FDR to adjust p-values. The Kruskal-Wallis test was used to compare the ΔRSCU for each codon across phage tRNA groups within each host genus to identify codons whose usage differs systematically among the tRNA groups. A simple effect size was computed from the Kruskal-Wallis statistic

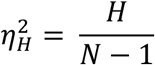

using the BH correction for FDR to adjust p-values across codons within a genus. A PCA was also run on ΔRSCU PERMANOVA data, with a matrix of (phages x codons). The scatterplot visualized is colored by lifestyle with shapes indicating the tRNA groups to visualize global clustering of phages based on these factors.

For comparison of ΔtAI values, preprocessing was used to generate variables amenable to modeling with the glmmTMB (50) package, which requires values to be strictly between 0 and 1. Before running statistical tests, the tAI data was rescaled, with any value ≤ 0 being increased to a very small value (1.00 x 10^-06^). A Wilcoxon signed-rank test with BH correction for FDR was used to determine the significance of the difference between a phage’s median tAI and the median tAI of its host. Data were tested against a null hypothesis of median ΔtAI = 0. The global ΔtAI beta mixed model uses the rescaled ΔtAI values per gene per phage using lifestyle, tRNA group, and host species as fixed effects and phage as the random effect (1|sample_id) to account for the fact that many genes come from the same phage and are not independent variables. This model was used to determine how lifestyle and tRNA group affect the relative adaptation of a phage to its host tRNA pool using the formula

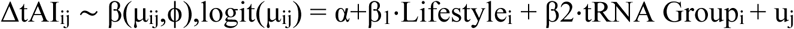

where u_j_ is a random intercept for each phage genome (sample_id). The group-level marginal means were extracted with emmeans (51) and the logit-transformed scale was back-transformed to tAI and ΔtAI values using the formula

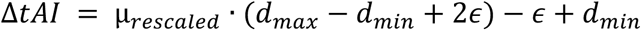

where µ_rescaled_ is the fitted mean from the beta regression, and d*_min_* and d*_max_* are the observed minimum and maximum ΔtAI values within the group. Like in the tAI calculation, ɛ is a small constant value (1.00 x 10^-06^) to prevent boundary issues. The same beta mixed model was used per host genus, including only the fixed effects of lifestyle and tRNA group along with the same random effect of phage.

All R scripts used in this analysis are available at github.com/nykkiross/phage-tRNAs.

## RESULTS AND DISCUSSION

### GC Content of Phages Compared to their Respective Hosts

The GC content of each phage was calculated and compared to the GC content of its host. The total GC content, as well as the codon position-specific GC contents – GC1, GC2, and GC3, representing codon positions 1, 2, and 3 respectively – were calculated for every coding sequence in a phage or host and the mean GC values for each genome were compared. Distributions of the GC content of each host genus and its infecting phages are visualized in **Figure 1**, and a summary of these values is provided in **Table 2**.

**Figure 1.**
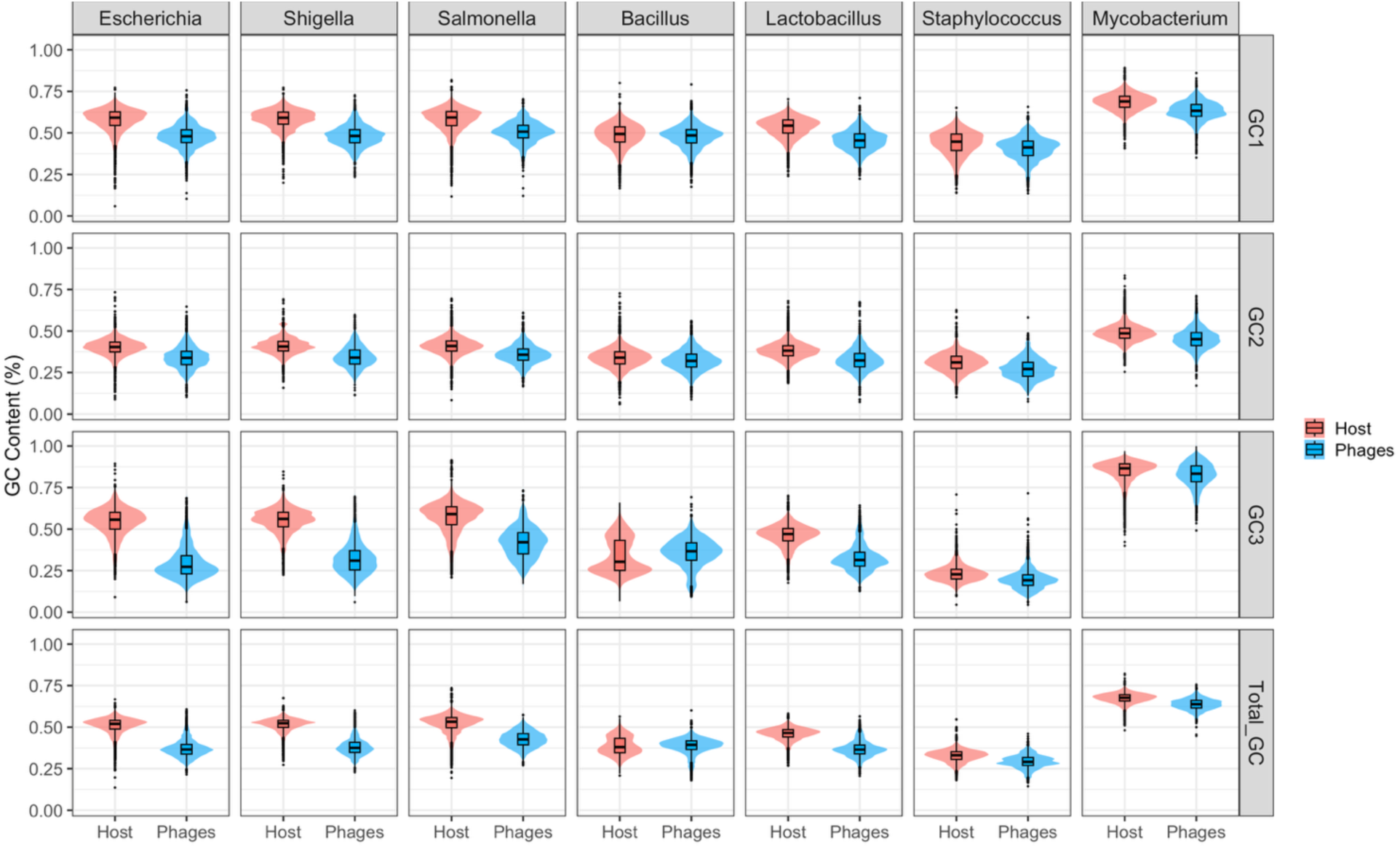
GC content of bacterial genera and their respective phages. The violin plot is faceted by host genus and GC type (GC1, GC2, GC3, Total GC). The GC content of all host genes was plotted compared to all phage genes for phages infecting that host genus. The distributions of these GC content values are plotted with a violin plot. Bacteria are shaded in red, phages in blue.

**Table 2.**
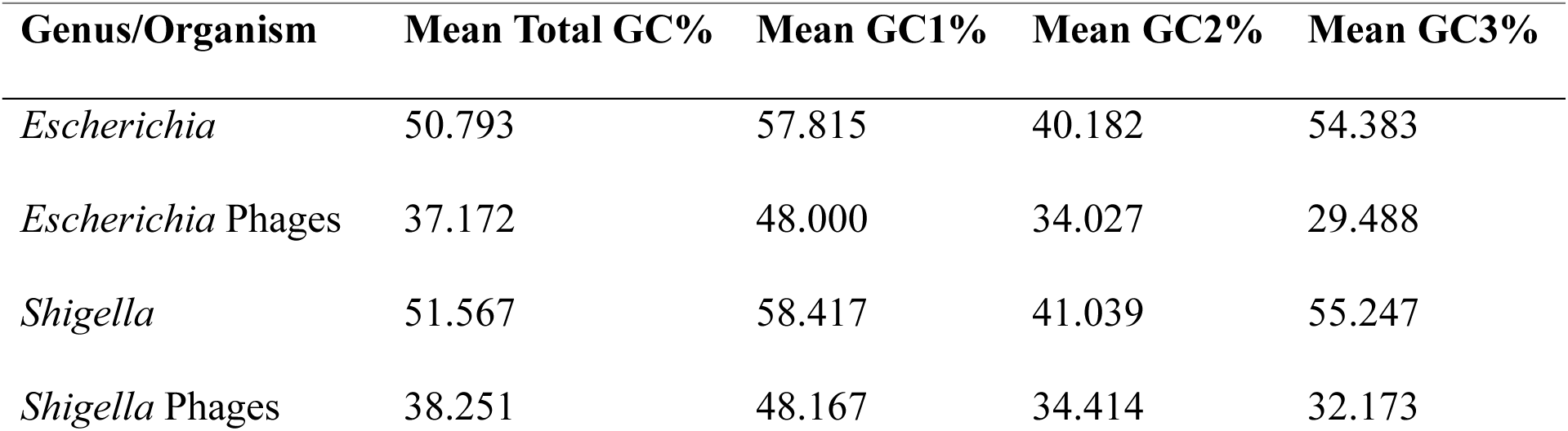
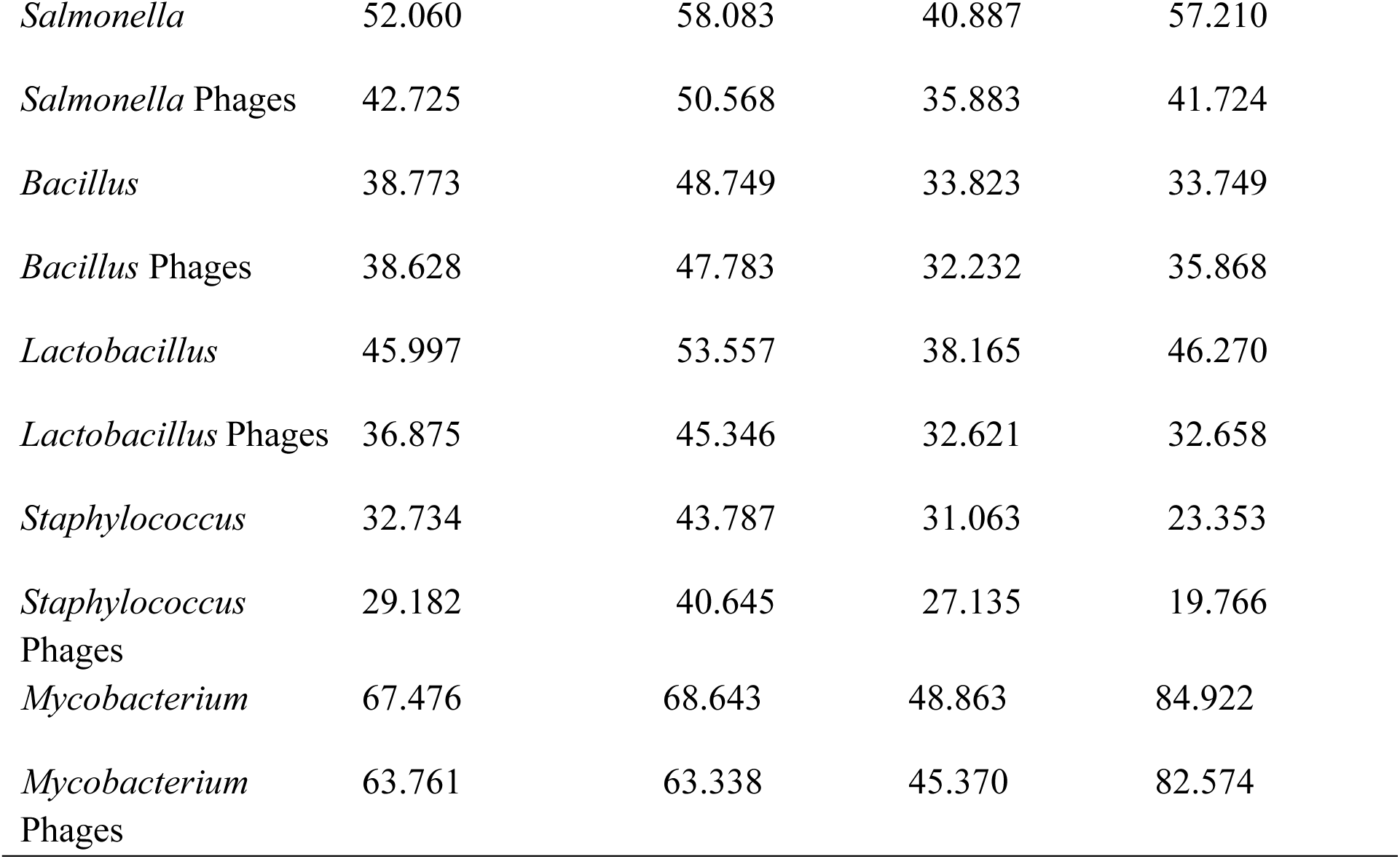
Summary of the mean GC content for phages and their host genera.

#### GC comparison of Gram-negative hosts and their phages

In general, the enteric bacteria investigated – *E. coli*, *S. flexneri*, *S. enterica* serovar Typhimurium, and *S. enterica* serovar Enteritidis – showed more significant differences between host and phage GC content. For example, the virulent *Escherichia* phage CAM-21 has a total GC of 34.71%, GC1 of 46.80%, GC2 of 32.95%, and GC3 of 24.42%, while its host *E. coli* has a total GC of 50.79% (p-value 3.74 x 10^-150^), GC1 of 57.81% (p-value 1.22 x 10^-98^), GC2 of 40.18% (p-value 4.87 x 10^-72^), and GC3 of 54.38% (p-value 1.29 x 10^-160^). The high significance of the difference in total GC as well as GC3 indicates codon bias has a role in genome composition. Virulent phages overall have more significant differences in GC content than their hosts compared to temperate phages, though some differences for temperate phages infecting enteric bacteria hosts still tend to be significant. For example, the temperate *Escherichia* phage lambda had a total GC of 50.08% (p-value 0.3895), GC1 of 56.26% (p-value 0.0307), GC2 of 42.66% (p-value 0.0006), and GC3 of 51.31% (p-value 0.0049). While the differences in total GC content between the phage lambda and its host *E. coli* are not significant (p-value > 0.05), the codon position-specific GC contents are. This indicates some level of genome composition differences, albeit not as strong as those seen in virulent phages.

Another example is with the host *S. flexneri*, which has a total GC of 51.57%, GC1 of 58.42%, GC2 of 41.04%, and GC3 of 55.25%. This contrasts significantly with the GC content of most *Shigella* phages, especially virulent phages, such as the virulent *Shigella* phage Sf24. This phage has a total GC of 34.51% (p-value 6.97 x 10^-163^), GC1 of 46.47% (p-value 1.35 x 10^-119^), GC2 of 32.57% (p-value 1.38 x 10^-89^), and GC3 of 24.49% (p-value 4.99 x 10^-166^). However, temperate *Shigella* phages show either much less significant or nonsignificant differences in GC content compared to the host. For example, the temperate phage SfII has a total GC of 50.16% (p-value 0.9346), GC1 of 56.14% (p-value 0.0698), GC2 of 42.43% (p-value 0.1151), and GC3 of 52.10% (p-value 0.2542). Likewise, the temperate *Shigella* phage SfV has a total GC of 50.80% (p-value 0.5000), GC1 of 57.60% (p-value 0.3410), GC2 of 42.69% (p-value 0.0272), and GC3 of 52.10% (p-value 0.0142). These phages were both induced from the *S. flexneri* genome as prophages (26,27), which may contribute to their more similar GC content.

The codon position-specific GC value which showed the least difference between host and phage across these bacterial hosts tended to be GC2, which is consistent with previous data (16), though the biological significance of this trend is unknown. Total GC content and GC3 content tend to have the most significant differences, indicating codon bias as a driving force behind genome composition differences. These trends are especially true for virulent phages.

#### GC comparison of Gram-positive hosts and their phages

Next, we investigated differences in GC content of phages infecting Gram-positive bacteria compared to their hosts. Phages infecting Gram-positive bacteria had slightly less significant differences in GC content compared to those seen in Gram-negative hosts, though these data are still highly significant across all comparisons. This could partially be due to the large number of temperate phages infecting the Gram-positive hosts included in this study, as these bacteria tended to have higher proportions of temperate phages compared to virulent phages.

While most of the host bacteria in this study have high GC content in their coding sequences, this is not the case for the *Bacillus* species (*B. cereus* and *B. subtilis*) analyzed here. *Bacillus* spp. are low-GC bacteria and thus show less significant differences between host and phage GC content for total and all codon position-specific GC contents. Interestingly, some of the phages infecting *Bacillus* spp. have significantly higher GC content than their hosts. This can be seen with the virulent *Bacillus* phage BCP78, which has a total GC of 39.50%, a GC1 of 48.88%, GC2 of 32.89%, and GC3 of 36.70%, compared to the host which has a total GC of 35.36% (p-value 4.78 x 10^-69^), GC1 of 46.92% (p-value 7.98 x 10^-05^), GC2 of 32.95% (p-value 0.5780), and GC3 of 26.21% (p-value 6.5 x 10^-110^). Consistent with the Gram-negative hosts, temperate phages tend to show less significant differences in GC content compared to their hosts. For example, the temperate *Bacillus* phage Gamma has a total GC of 34.84% (p-value 0.2592), GC1 of 45.84% (p-value 0.2095), GC2 of 31.31% (p-value of 0.0800), and GC3 of 27.36% (p-value 0.0907). The host *Staphylococcus aureus* is also a low-GC host and phages infecting *Staphylococcus* spp. show similar patterns in GC content when compared to the host as those seen in *Bacillus* phages.

While most phages in this analysis had low GC content, the *Mycobacterium* phages have comparatively high GC values, which may reflect the high GC content of the host *M. smegmatis*. Like the other Gram-positive hosts, there is a less significant difference in GC content between phages and the *Mycobacterium* host compared to the differences seen in the Gram-negative bacteria. Whether a phage is temperate or virulent seems to have the opposite impact on GC content, with temperate phages showing more significant differences compared to the host than virulent phages. For example, the host *M. smegmatis* has a total GC of 67.48%, GC1 of 68.64%, GC2 of 48.86%, and GC3 of 84.92%, while the virulent phage Virapocalypse has a total GC of 66.49% (p-value 0.0001), a GC1 of 66.11% (p-value 3.73 x 10^-07^), GC2 of 50.51% (p-value 0.0050), and GC3 of 82.84% (p-value 5.89 x 10^-07^). Conversely, the temperate phage Iwokeuplikedis shows more significant differences in its GC content, with a total GC of 62.63% (p-value 2.62 x 10^-41^), GC1 of 62.05% (p-value 1.62 x 10^-27^), GC2 of 43.67% (p-value 1.11 x 10^-15^), and GC3 of 82.18% (p-value 3.20 x 10^-09^). Like the other Gram-positive hosts, some phages show higher GC content than their host.

Like the temperate phages of Gram-negative hosts, the Gram-positive temperate phages tend to show less significant differences in GC content than virulent phages for total and all codon position-specific GC values, with GC2 showing the least significant difference for most phages. The exception is seen in the *Mycobacterium* phages, which show the least significant differences for GC3. With the least significant differences occurring at the wobble base, this argues against GC content driving codon usage differences. The pattern of GC content for total GC and GC3 across Gram-positive hosts is less clear, indicating codon bias may have a lesser impact on the genome composition of phages infecting these hosts.

### ΔGC Comparison of Phages According to tRNA Group

The ΔGC content for total GC, GC1, GC2, and GC3 were calculated for each phage by subtracting the mean host GC content from the mean phage GC content for each phage and each host. These data were then used to compare phages both within host genus and globally across all host genera. ΔGC was used rather than GC for these analyses due to differences in genome composition between the hosts. A heatmap depicting the ΔGC data for all GC types in the tRNA group comparisons for each host genus is available in **Figure 2**.

**Figure 2.**
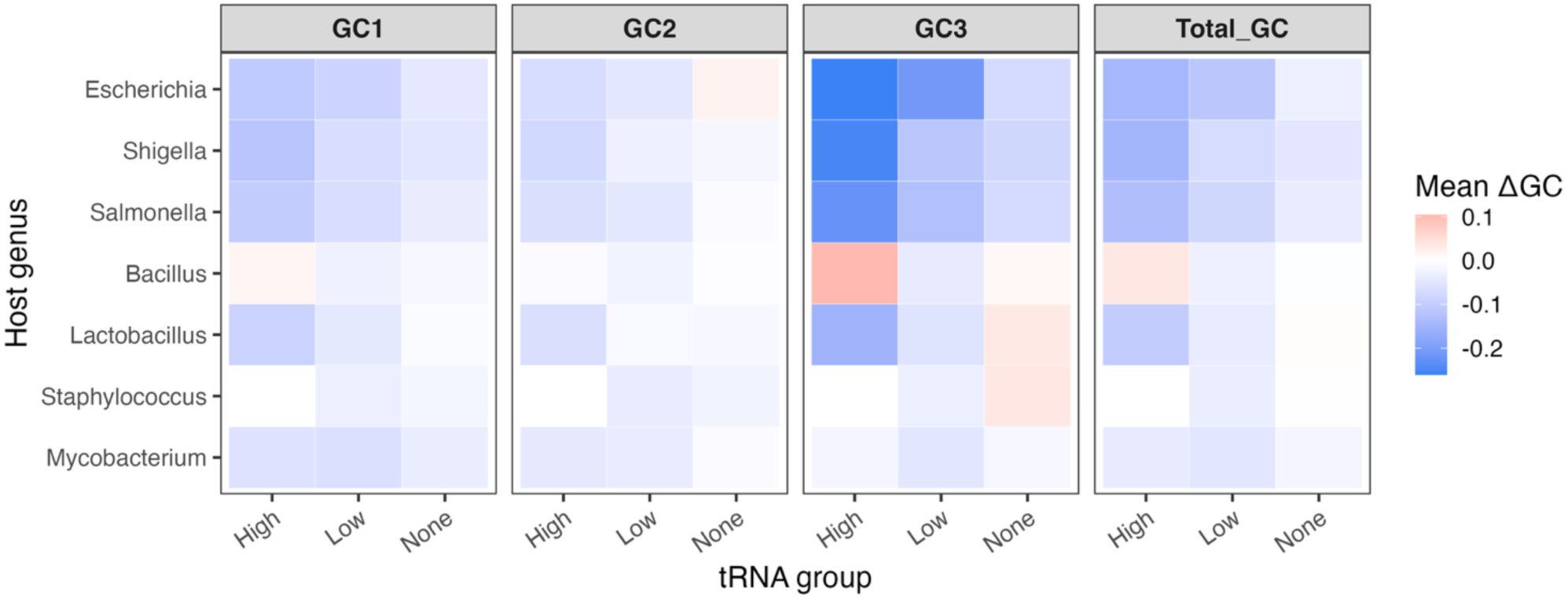
ΔGC by tRNA group. The heatmap is faceted by GC type (GC1, GC2, GC3, Total GC) and shows the mean ΔGC for each GC type for each tRNA group (High, Low, None). Lower values indicate significantly lower GC content in the phages, while higher values indicate similar or higher GC content. Lower values are blue, and higher values are pink.

#### ΔGC comparison of tRNA groups within each genus

For this analysis, the ΔGC for total GC, GC1, GC2, and GC3 were compared across phage tRNA groups. The tRNA groups consist of High (> 5 tRNAs), Low (1 – 5 tRNAs), and None (0 tRNAs). The ΔGC content was compared between the High vs. Low groups, the High vs. None groups, the Low vs. None groups, and lastly the High and Low groups were combined and compared to the None group, in order to determine if the number of tRNAs a phage encodes and if encoding tRNAs in general affect how different a phage’s GC content is from its host.

In *Escherichia* phages, there is not a significant difference in ΔGC content between phages with high numbers of tRNAs and those with low numbers of tRNAs, and there is no significant difference between phages with low numbers of tRNAs and those carrying none. However, a significant difference in ΔGC content is seen between phages with high numbers of tRNAs and those with none for all GC types (p-values range from 0.0019 to 0.0025), with higher numbers of tRNAs associated with larger differences in GC content from the host. Having tRNAs in general results in a significantly different ΔGC content compared to carrying none for all GC types as well (p-values range from 3.62 x 10^-04^ to 0.0015).

Conversely, *Shigella* phages only showed nonsignificant differences in ΔGC content between the Low vs. None groups. The High tRNA group had significantly different ΔGC content compared to both the Low group (p-values range from 0.0095 to 0.0131) and the None group (p-values range from 0.0037 to 0.0050) for all GC types with the exception of ΔGC2. This suggests that *Shigella* phages with many tRNAs have significantly larger differences in GC content than their host compared to low or no tRNAs. Likewise, having tRNAs in general resulted in significantly larger ΔGC values compared to having no tRNAs (p-values range from 0.0028 to 0.0392). *Salmonella* phages exhibited significantly different ΔGC content for the High vs. Low comparison for all GC types, with phages in the High tRNA group exhibiting larger absolute ΔGC values (p-value 0.0106 for all GC types). Conversely, the High vs. None group comparison and the Low vs. None group comparison did not show significant differences in these values. In looking at phages with tRNAs in general, regardless of number, the ΔGC content for all GC types is also significantly different compared to those without tRNAs (p-values range from 0.0098 to 0.0168). From this analysis, phages of Gram-negative bacteria with high numbers of tRNAs exhibit larger ΔGC values compared to their respective hosts, suggesting that the tRNAs may compensate for genome compositional differences between phage and host.

Interestingly, only the High vs. Low groups for *Bacillus* phages show significant differences in ΔGC content for all GC types, with the High tRNA group showing a significantly smaller difference from the host GC content compared to phages in the Low tRNA group (p-values range from 0.0056 to 0.0397). Significant differences are not seen in the High vs. None, Low vs. None, nor combined High + Low vs. None comparisons. For *Staphylococcus* phages, only the Low vs. None groups could be compared, as no phage had more than 4 tRNA genes. Comparing these two groups still revealed significant differences in ΔGC content, with larger differences in GC content compared to the host seen in phages carrying tRNAs (p-values range from 0.0189 to 0.0336), with the exception of ΔGC1. Phages infecting *Lactobacillus* spp. show significant differences in ΔGC content only when comparing the High + Low combined group with the None group for most GC types (p-value 0.0307); however, ΔGC2 showed no significant differences. In *Lactobacillus* phages, having tRNAs in general is associated with a significantly larger difference in GC content from the host than seen in phages without tRNAs. Conversely, in the High vs. Low group comparison the only significant difference was seen in ΔGC2, with the High group having a significantly higher absolute ΔGC2 (p-value 0.0081).

The only significant difference in ΔGC content seen in phages infecting *Mycobacterium* was seen for total ΔGC in the High + Low combined vs. None group comparison (p-value 0.0431), with phages containing tRNAs in general having larger differences in GC content compared to their host than phages without tRNAs. Since phages infecting Gram-positive hosts seem to match their GC content more with their hosts’, this result is not surprising and provides less support for tRNAs compensating for differences in GC content in these phages.

#### Global ΔGC comparison of tRNA groups across genera

To determine the impact of the number of tRNA genes carried by a phage on the ΔGC content, a global comparison was performed, combining the groups of phages from each host genus into High, Low, and None groups. When all ΔGC data is pooled together, there are significant differences in ΔGC content for all GC types for all comparisons except the Low vs. None comparison, where only ΔGC2 was significantly different, being larger in the Low group (p-value 4.55 x 10^-04^). There were significant differences in ΔGC content for all GC types for the High vs. Low comparison (p-values range from 3.77 x 10^-08^ to 1.01 x 10^-05^), High vs. None comparison (p-values range from 8.59 x 10^-16^ to 2.00 x 10^-08^) and the combined High + Low vs. None comparison (p-values range from 7.60 x 10^-11^ to 2.45 x 10^-05^). The High groups consistently have larger absolute ΔGC values for all GC types, indicating a larger difference in GC content of these phages compared to their hosts. On a global scale, the number of tRNAs a phage encodes seems to be related to how different a phage’s GC content is from its host, with higher numbers of tRNAs associated with larger differences in genome composition.

### ΔGC Comparison of Phages According to Lifestyle

The ΔGC content data generated for the tRNA group comparisons was then used to determine if the lifestyle of a phage impacts how much the GC content of a phage differs from that of its host. Phages were grouped into either Virulent or Temperate groups, then compared both within each host genus and globally across genera. A heatmap depicting the ΔGC data for all GC types in the lifestyle comparisons for each host genus can be found in **Figure 3**.

**Figure 3.**
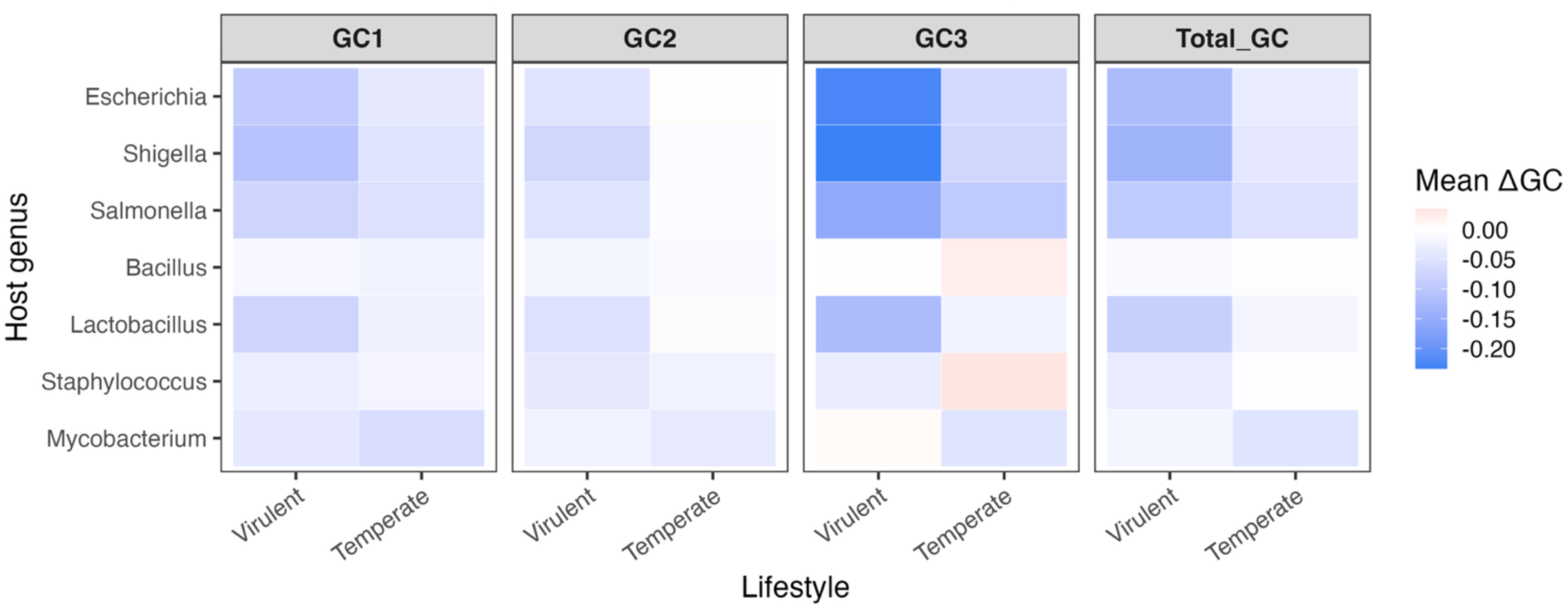
ΔGC by Phage Lifestyle. The heatmap is faceted by GC type (GC1, GC2, GC3, Total GC) and shows the mean ΔGC for each GC type for both lifestyles (Virulent or Temperate). Lower values indicate significantly lower GC content in the phages, while higher values indicate similar or higher GC content. Lower values are blue, and higher values are pink.

#### ΔGC comparison of lifestyle within each genus

There are significant differences in ΔGC content for all GC types between virulent and temperate phages for *Escherichia* phages (p-values range from 9.88 x 10^-04^ to 0.0227) and *Shigella* phages (p-values range from 2.97 x 10^-05^ to 2.82 x 10^-04^), with virulent phages having significantly larger absolute ΔGC for all comparisons. This was not the case for *Salmonella* phages, which only showed a significant difference in ΔGC2. Virulent phages had significantly higher ΔGC2 compared to temperate phages (p-value 0.0346). It must be noted that the virulent and temperate groups of the *Salmonella* phages are imbalanced, as only two temperate phages met the criteria for the analysis. This was taken into account during all statistical analyses.

There were no significant differences in ΔGC content for virulent and temperate *Bacillus* phages for any GC type, but *Mycobacterium* and *Staphylococcus* phages showed significant differences in ΔGC content for all GC types except ΔGC2 (p-values range from 1.72 x 10^-05^ to 0.0042 in the former and from 0.0151 to 0.0303 in the latter). Interestingly, these phages show the opposite trend compared to phages infecting Gram-negative hosts, with temperate phages having larger absolute ΔGC values than virulent phages. Phages infecting *Lactobacillus* were the only phages infecting Gram-positive bacteria with significant differences in the ΔGC content of all GC types with no exceptions (p-values range from 0.0151 to 0.0319). The *Lactobacillus* phages showed ΔGC patterns similar to phages infecting Gram-negative hosts, with virulent phages having larger absolute ΔGC for all GC types compared to temperate phages.

Whether the lifestyle of a phage impacts how much its GC content differs from that of its host seems to be host-dependent, with phages infecting Gram-negative hosts consistently having more significant differences. Whether the virulent phages or temperate phages differ more in GC content also seems to be host-dependent. Virulent phages infecting Gram-negative hosts, along with virulent phages of the Gram-positive host *Lactobacillus*, have larger ΔGC values compared to phages infecting the other Gram-positive hosts or compared to temperate phages.

#### Global ΔGC comparison of lifestyle across genera

To assess the effect of a phage’s lifestyle on ΔGC content globally, virulent phages from all hosts and temperate phages from all hosts were combined and compared. The ΔGC content was highly significant (p-values range from 4.68 x 10^-06^ to 0.0028) for all GC types when comparing virulent and temperate phages across all host genera. This indicates that lifestyle has a significant effect on the ΔGC content of a phage, with virulent phages differing significantly more from host GC content compared to temperate phages. Since this was observed across all genera, it is likely reflective of the long-term relationship between temperate phages and their hosts compared to virulent phages, rather than host-specific effects.

### Effective Number of Codons (EnC) of Phages Compared to their Respective Hosts

The effective number of codons (EnC) is a way of measuring the number of unique codons in a gene (42). This index returns a single value per gene reflecting the quantity of unique codons in that gene. Values range from 20 (highly biased; only one codon used per amino acid) to 61 (low bias; every codon for every amino acid is used). The effective number of codons itself doesn’t indicate which direction codon bias is in, since it does not address the GC content of a gene or the frequency of the codons which are used in the gene; however, it can be used to estimate the impact of selection and genetic drift on codon usage. Higher EnC values, reflecting lower levels of bias, indicate that codon usage differences are because of genetic drift. Conversely, lower EnC values are associated with higher levels of codon bias in a gene, indicating selective pressures on codon usage. Based on the results described here, nearly all phages show larger EnC values compared to their hosts, regardless of host genus. This indicates a lower level of codon bias in phages compared to bacteria. A summary of the EnC data for hosts and phages can be found in **Table 3**.

**Table 3.** Summary of median EnC values in phages and their hosts.

| Host Genus | N Host | N Phage | Median EnC<br>Host | Median EnC<br>Phage | Std dev<br>EnC |
| --- | --- | --- | --- | --- | --- |
| *Escherichia* | 1 | 24 | 38.621 | 40.594 | 2.309 |
| *Shigella* | 1 | 24 | 39.323 | 41.969 | 2.173 |
| *Salmonella* | 2 | 22 | 38.480 | 42.855 | 1.556 |
| *Bacillus* | 2 | 23 | 39.235 | 44.705 | 2.543 |
| *Lactobacillus* | 3 | 17 | 41.093 | 41.165 | 2.382 |
| *Staphylococcus* | 1 | 23 | 36.318 | 38.656 | 2.474 |
| *Mycobacterium* | 1 | 21 | 31.282 | 36.459 | 2.409 |

#### EnC comparison of Gram-negative hosts and their phages

While GC content showed fairly consistent patterns of significance when comparing phages to their hosts, this was not the case with EnC comparisons. The most variation was seen in the *Escherichia* phages, with 11 phages showing highly significant differences in EnC compared to *E. coli* (p-values range from 8.60 x 10^-48^ to 5.93 x 10^-04^) and 13 phages showing nonsignificant differences in EnC. Regardless of significance, phages consistently show higher EnC values compared to their hosts. The lifestyle and the number of tRNA genes carried by a phage do not seem to have an impact on whether a phage’s EnC is significantly different from its host’s; rather, the classification of a phage does seem to play a role. Specifically, the *Straboviridae* phages consistently showed nonsignificant differences in EnC compared to the host, indicating higher levels of codon bias in these phages compared to other *Escherichia* phages.

The *Shigella* phages also had variation in the significance of EnC differences between each phage and its host. Significant differences were seen mainly in *Ounavirinae* phages, but the temperate phages SfII and SfV, two myoviruses, also showed significant differences in EnC values compared to *S. flexneri* (p-values range from 2.33 x 10^-22^ to 2.04 x 10^-04^). Nonsignificant differences in EnC were specifically seen only in the *Straboviridae* phages and the *Schitoviridae* phages, but taxonomy-specific patterns were not seen when computing any other index. Whether a phage is temperate or virulent and the number of tRNA genes carried by a phage do not seem to affect the significance of the difference in EnC values, though. Conversely, every *Salmonella* phage showed significant differences in EnC from its host (p-values range from 3.14 x 10^-57^ to 1.17 x 10^-06^), regardless of number of tRNA genes, lifestyle, or taxonomic classification. Overall, phages of *Escherichia*, *Shigella*, and *Salmonella* consistently have higher EnC values compared to their hosts, indicating lower levels of codon bias.

#### EnC comparison of Gram-positive hosts and their phages

Nearly all of the *Bacillus* phages showed highly significant differences in EnC values compared to both *B. cereus* and *B. subtilis* bacteria (p-values range from 2.43 x 10^-51^ to 4.35 x 10^-04^), with the exception of the *B. subtilis* phage φ29. Significantly different EnC values were seen in both temperate and virulent phages, regardless of the tRNA gene number carried by a phage. Likewise, nearly every phage infecting *Staphylococcus* showed highly significant differences in EnC compared to *S. aureus* (p-values range from 3.01 x 10^-11^ to 0.0052), with the exception of the temperate phage SAP-2 and the virulent phage Sb-1. *Mycobacterium* phages showed highly significant differences in EnC compared to the host (p-values range from 8.88 x 10^-37^ to 0.0183), with the exception of virulent phage Phabba, which carries the highest number of tRNAs of any phage in this study with 37 tRNA genes. Similar to *Bacillus*, lifestyle and number of tRNA genes carried by a phage do not seem to affect the significance of the difference in EnC values in *Staphylococcus* or *Mycobacterium* phages.

It may be notable to point out that, while nearly all phages in these analyses showed higher EnC values compared to their hosts, the *Staphylococcus* phages Machias, S6, and SAP-2 have lower EnC values than their host. For Machias and S6, this difference is significant (p-values 4.37 x 10^-07^ and 5.95 x 10^-04^, respectively). The only other phage which showed a lower EnC than that of its host is the *Mycobacterium* phage Phabba, but this difference was not significant.

The *Lactobacillus* phages were the exception to this trend in Gram-positive hosts, with only 8 phages showing significant differences in EnC compared to their host (p-values range from 2.49 x 10^-11^ to 0.0031). The remaining phages of the virulent *Herelleviridae* family showed nonsignificant differences. While phage classification had a specific impact across more than one Gram-negative host, *Lactobacillus* phages were the only phages infecting Gram-positive hosts in which phage classification had an impact. For example, most *Staphylococcus* phages also belong to the *Herelleviridae* family, but they all show significant differences in EnC compared to the host. The number of tRNA genes carried by a phage also does not seem to impact the significance of the difference in EnC values of *Lactobacillus* phages compared to their hosts; conversely, lifestyle may have an effect, with virulent phages having EnC values closer to that of their host.

### ΔEnC Comparison of Phages According to tRNA Group

Next, the ΔEnC was calculated for each phage by subtracting the mean host EnC from the mean phage EnC. These data were then used to compare phages based on the number of tRNAs encoded in their genomes, both within a host genus and globally across host genera. ΔEnC was used for this analysis rather than EnC to account for differences in host EnC values.

#### ΔEnC comparison of tRNA groups within each genus

As with the ΔGC comparisons, phages were grouped per host genus into one of three groups according to number of tRNA genes encoded in their genomes – High (> 5 tRNAs), Low (1 – 5 tRNAs), or None (0 tRNAs). Within each host, the ΔEnC were compared between the High vs. Low groups, High vs. None groups, and Low vs. None groups. This allowed us to determine if the number of tRNA genes in a phage genome has an impact on how different its EnC is from its respective host. A combined High + Low group was also compared to the None group to determine if having tRNAs in general affects how different a phage’s EnC is from its host’s.

In general, the number of tRNA genes did not have an effect on the differences between host and phage EnC values, regardless of the tRNA groups being compared. There were only a few exceptions. First, when comparing *Salmonella* phages with and without tRNAs, it was found that phages with tRNAs have significantly smaller ΔEnC values (p-value 0.0353). While the Low vs. None comparison did not show significant differences, the High group had a significantly smaller ΔEnC (median 3.261) than the None group (median 7.324; p-value 0.0244). This indicates that *Salmonella* phages with high numbers of tRNAs have EnC values closer to those of the host compared to phages with no tRNAs, and thus exhibit stronger codon bias. Additionally, the *Lactobacillus* High vs. Low comparison (p-value 0.0425) and High vs. None comparison (p-value 0.0425) were also significant, with the High group having a median ΔEnC of 0.361, the Low group having a median ΔEnC of 4.149, and the None group having a median ΔEnC of 4.386. These results also indicate that phages carrying high numbers of tRNAs differ less in their EnC values from the host compared to those with low or no tRNAs, supporting the idea that *Lactobacillus* phages with tRNAs show stronger codon usage bias. In other hosts, the number of tRNAs in a phage genome appears to have very little impact on how different a phage’s EnC is compared to that of its host.

#### ΔEnC comparison of tRNA groups across genera

When compared globally, different patterns were seen. A global comparison of the High vs. Low groups resulted in a significant difference in their ΔEnC values with the High group having a median ΔEnC of 2.976 and the Low group having a median of 4.117 (p-value 0.0026). Additionally, the comparison of ΔEnC values between the High vs. None groups was also highly significant, with the None group having a median of 4.551 (p-value 6.71 x 10^-04^). Finally, when the High and Low groups were combined and compared to the None group, the ΔEnC was significantly different (p-value 0.0094). However, the difference in ΔEnC between the Low vs. None groups globally showed no significant difference. These data taken together indicate that on a global scale, having high numbers of tRNAs is associated with smaller differences in EnC compared to the host, suggesting a higher level of codon bias in the genomes of phages with high numbers of tRNAs compared to those with low or no tRNAs.

### ΔEnC Comparison of Phages According to Lifestyle

The ΔEnC values for each phage were then used to compare phages by lifestyle to determine if being virulent or temperate results in EnC values more or less different from that of the host. These analyses were performed both within each host genus and globally across host genera.

#### ΔEnC comparison of lifestyle within each genus

No Gram-negative host showed significant differences in ΔEnC when comparing phage lifestyles, though the ΔEnC values for virulent phages were consistently lower than those for temperate phages. Only two host genera were infected by phages with significant differences in ΔEnC when the phages are temperate versus virulent. The *Mycobacterium* phages show significantly larger ΔEnC for temperate phages (median 5.480) compared to virulent phages (median 3.028, p-value 0.0037). Phages infecting *Lactobacillus* also showed significant differences in ΔEnC, with temperate phages having significantly larger ΔEnC values than virulent phages (median 4.249 vs median 0.459, p-value 0.0404). In both of these cases, the results suggest that the genomes of virulent *Mycobacterium* and *Lactobacillus* phages have significantly stronger codon bias than the temperate phages.

While lifestyle does not seem to have a significant impact on the distance between host and phage EnC values for most hosts, virulent phages consistently had smaller ΔEnC values, with host EnC values lower than the phages’ EnC values in the phage to host comparisons. With virulent phages having EnC values closer to those of their hosts, this indicates higher levels of codon bias. Conversely, the temperate phages have larger EnC values and a larger ΔEnC when compared to their hosts, indicating these phages have lower levels of codon bias.

#### ΔEnC comparison of lifestyle globally across genera

When comparing the ΔEnC of temperate and virulent phages by genus, significant differences were not seen in every host; however, when phages are pooled and analyzed globally, the difference in ΔEnC between temperate and virulent phages was highly significant (p-value 7.52 x 10^-09^). Temperate phages globally had significantly larger ΔEnC than virulent phages (median 5.44 vs. 3.00, respectively). Overall, the results indicate that across all genera investigated, virulent phages show smaller differences in EnC values compared to their hosts than temperate phages. This leads to the conclusion that virulent phages have higher levels of codon bias, as the hosts consistently had lower EnC values – associated with higher codon bias – than the phages that infect them.

### Correlation Analysis of EnC to GC3

The correlation between the EnC value and GC3 value for a gene can provide insight into the impact of selective pressure on codon usage. Results were calculated using Spearman’s correlation. Very weak correlations have rho values < 0.10, weak correlations 0.10 - 0.30, moderate correlations 0.30 - 0.50, and strong correlations > 0.50. Stronger correlations indicate selection is responsible for the codon usage biases seen in a gene or genome, while weaker correlations suggest that genetic drift is the reason for the codon usage biases. The results can then be visualized using an Nc plot, and the impact of selective pressure versus that of genetic drift can be estimated. Values that fall on or above Wright’s expected Nc curve indicate genetic drift is responsible for codon bias in that gene (45). Values that fall below the Nc curve indicate selective pressure is present and has an impact on codon bias in a gene.

#### Correlation between EnC and GC3 for phages according to host genus

For most phages, the Spearman’s correlation between EnC and GC3 was positive, meaning that as EnC increases, so does GC3 content. Subsequently, phages with low GC3 content should theoretically exhibit high levels of codon bias compared to those with high GC3 content. For the majority of hosts, this correlation was weak to moderate. In terms of Gram-negative genera, both *Escherichia* and *Shigella* phages had a moderate, positive correlation between EnC and GC3 (rho values 0.355 and 0.312, respectively). These relationships were also highly significant (p-values 8.65 x 10^-106^ and 2.36 x 10^-70^, respectively). *Salmonella* phages, however, did not appear to show a correlation between EnC and GC3 (rho value of 0.082, p-value 3.33 x 10^-06^).

For Gram-positive genera, phages infecting *Bacillus* and *Lactobacillus* hosts also showed moderate positive and highly significant relationships between EnC and GC3 (respective rho values 0.376 and 0.403, p-values 3.34 x 10^-125^ and 4.29 x 10^-74^). The phages infecting *Staphylococcus* show the strongest positive correlation of the hosts mentioned thus far, with a rho value of 0.418 (p-value 3.26 x 10^-142^). Finally, *Mycobacterium* phages were the exception to the patterns seen in phages infecting other host genera. *Mycobacterium* phages show a highly significant but strong negative correlation between EnC and GC3 (rho value -0.641, p-value 2.75 x 10^-277^). These results indicate that for *Mycobacterium* phages specifically, GC3 values decrease as EnC values increase, so phages with higher codon bias have higher GC3 values. These results align with the observed high GC content of the host and the phages infecting *Mycobacterium*.

Overall, there was a moderate relationship between EnC and GC3 in the majority of phages within each host genus. This supports some impact of GC3 on codon bias, but other factors like genetic drift or adaptation to a host genome may also affect the level of codon bias in a phage genome. For nearly all hosts, phages show a positive relationship between EnC and GC3, indicating that as codon bias increases, both EnC values and GC3 content decreases. This results in stronger codon biases in the AT-rich phage genomes. The exception to this is the *Mycobacterium* phages, which show a negative correlation likely due to the high GC content of the host and phage genomes. The relationship between EnC and GC3 was also the strongest for *Mycobacterium* phages, pointing to a larger role of selective pressure in the codon usage of these phages. Nc plots showing the comparison between phages and their hosts are shown in **Figure 4**.

**Figure 4.**
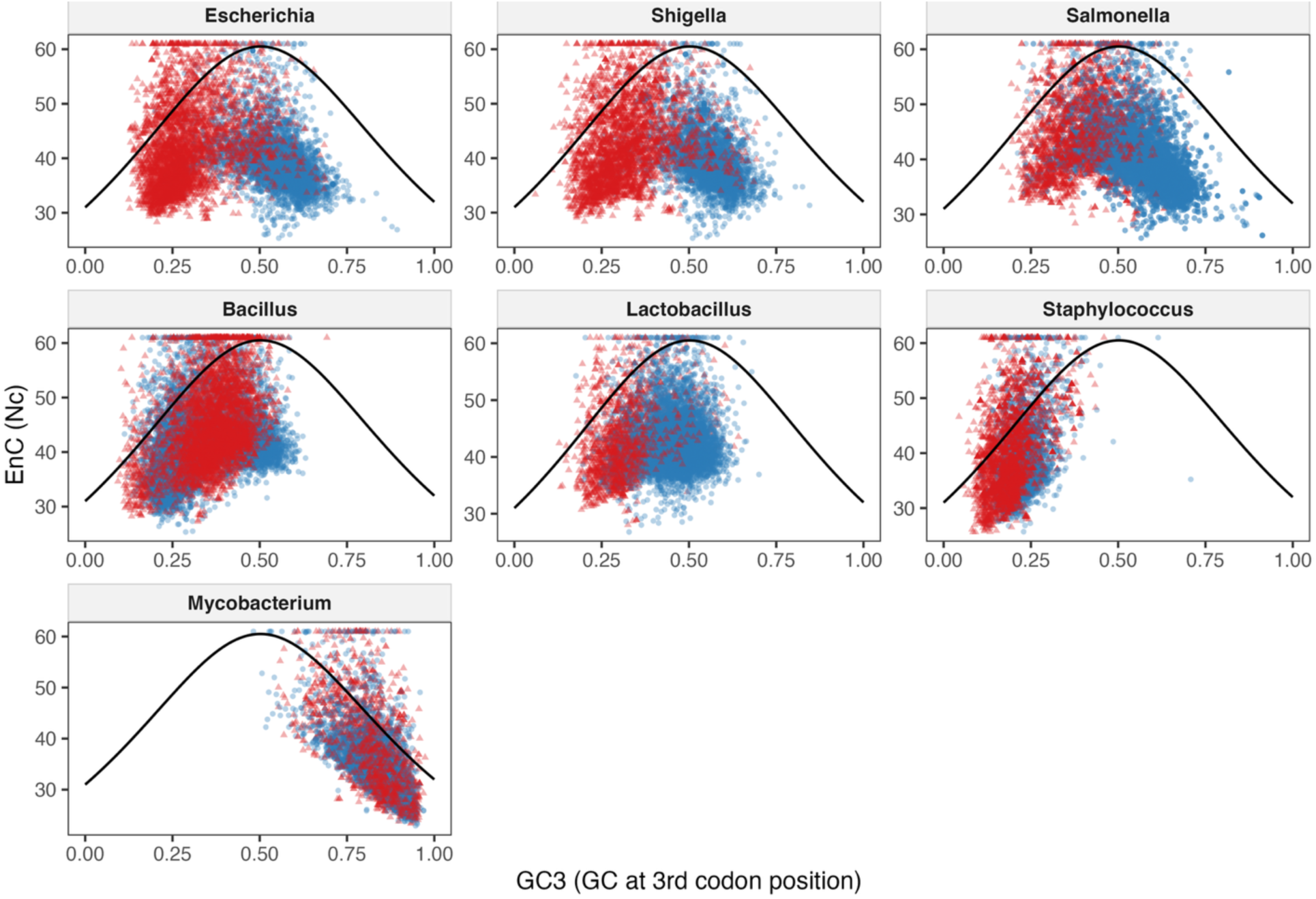
Nc plot of bacterial genera and their respective phages. The Nc plot depicts the relationship between EnC and GC3 for all phage and respective host genes. The plots are faceted by genus, with bacteria represented by blue circles and phages represented by red triangles.

#### Correlation between EnC and GC3 by tRNA group according to host genus

We next examined the differences in the EnC and GC3 correlations for each of the tRNA groups, which can provide insight into whether the number of tRNAs carried by a phage affects the strength and significance of this relationship. Nc plots showing the comparison between phages based on tRNA group per host genus are shown in **Figure 5**, with Gram-negative bacteria in the top row. In *Escherichia* phages belonging to the High and Low groups, the relationship between EnC and GC3 is positive and moderate (rho value 0.353 and 0.350, respectively) and highly significant for both the High group (p-value 8.32 x 10^-85^) and Low group (4.46 x 10^-15^). In the None group, a weak negative correlation between EnC and GC3 was found (rho value -0.143), but this correlation was not significant. Conversely, *Shigella* phages showed a weak positive relationship between EnC and GC3 for the High group only (rho value 0.292), which was highly significant (p-value 5.08 x 10^-47^). The Low group had a very weak, nonsignificant correlation between EnC and GC3 (rho value 0.090), while the None group had a very weak, nonsignificant negative correlation (rho value - 0.014). Similar to *Shigella* phages, *Salmonella* phages in the High group showed the strongest correlation between EnC and GC3, with a moderate positive correlation (rho value 0.324, p-value 6.51 x 10^-24^). The Low and None groups both had very weak negative correlations (rho values - 0.065 and -0.099 respectively), but only the correlation seen in the Low group was significant (p-value 0.003). Combined, these results show that for Gram-negative bacteria and their phages, the number of tRNAs encoded by a phage seems to increase the strength of the relationship between EnC and GC3. This indicates that high numbers of tRNAs may be associated with stronger selective pressures on codon bias in the AT-rich phage genomes.

**Figure 5.**
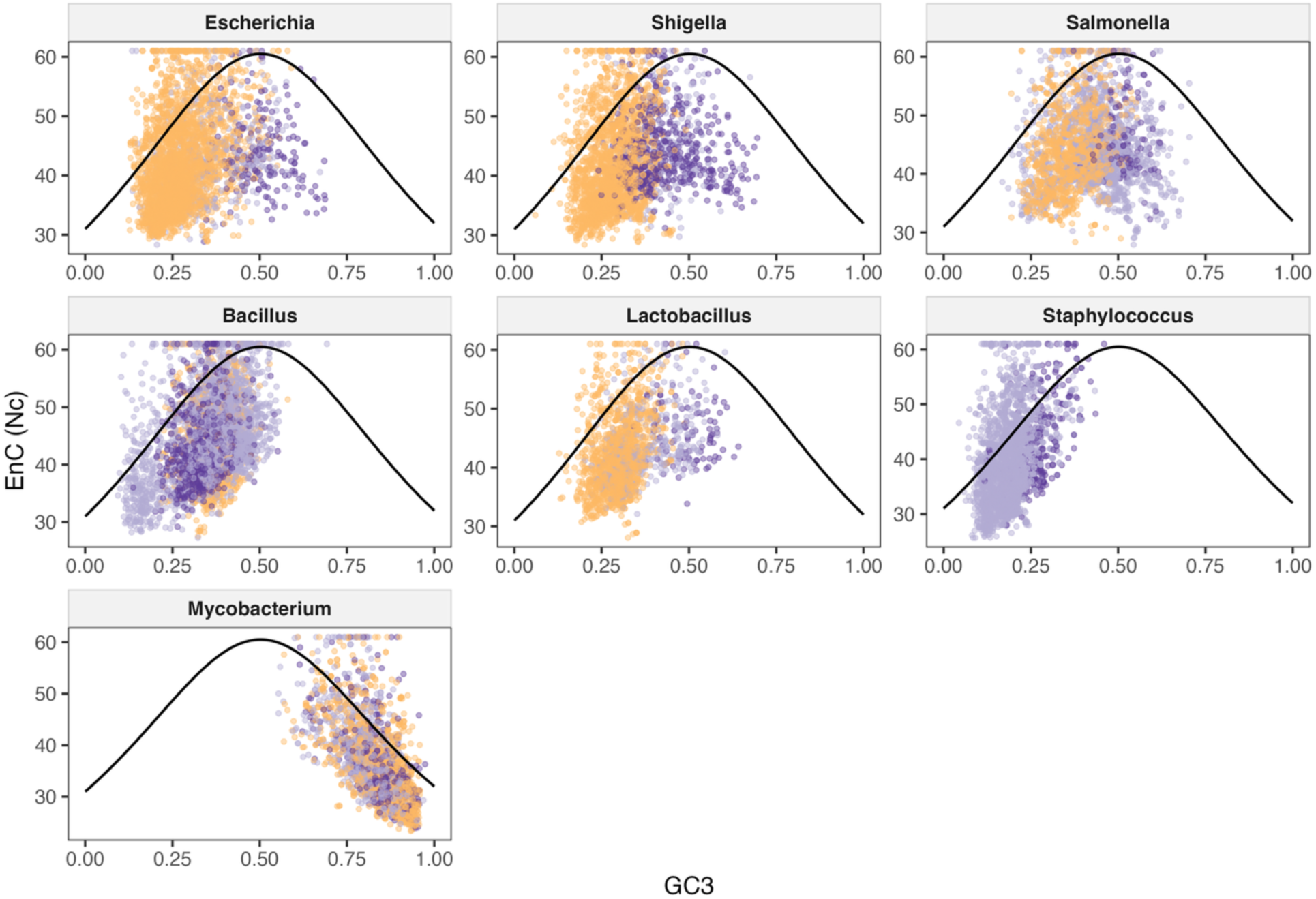
Nc plot of phages according to tRNA group. The Nc plots are faceted by host genus and depict the relationship between EnC and GC3 for each phage in each of the three tRNA groups (High, Low, None). The plot shows datapoints from all genes in all phages infecting that genus. High tRNA phages are colored in yellow, Low tRNA phages are colored in light purple, and None tRNA phages are colored in dark purple.

Unlike the patterns seen for Gram-negative hosts, phages infecting Gram-positive bacteria generally show stronger correlations when there are few or no tRNAs in the genome. For example, in *Bacillus* phages, there is a moderate positive correlation between EnC and GC3 for both the Low and None groups (rho values 0.423 and 0.304, respectively), both of which are highly significant (p-values 5.95 x 10^-97^ and 8.24 x 10^-12^, respectively). The High group showed a weak but significant positive correlation (rho value 0.185, p-value 7.00 x 10^-09^). *Staphylococcus* phages also showed moderate positive correlations between EnC and GC3 for both the Low and None tRNA groups (rho values 0.398 and 0.495, respectively) which were both highly significant (p-values 2.21 x 10^-118^ and 5.61 x 10^-16^, respectively). For *Staphylococcus* phages, the High group could not be compared, as none of the phages infecting *Staphylococcus* investigated had more than 4 tRNAs. *Lactobacillus* phages show a pattern more similar to *Escherichia* phages than other Gram-positive hosts, with the High and Low groups showing moderate positive correlations (rho values 0.330 and 0.461, respectively), both of which were highly significant (p-values 5.69 x 10^-38^ and 1.53 x 10^-18^, respectively). The None group showed a weak negative correlation (rho value -0.123) that was nonsignificant. Therefore, for most of these phages, lower numbers of tRNAs seem to be associated with a stronger impact of selective pressures on codon bias. The Nc plots for these Gram-positive bacteria are in the middle row of **Figure 5**.

The *Mycobacterium* phages once again are unique in that all tRNA groups have a strong negative correlation between EnC and GC3. As shown in the bottom row of **Figure 5**, the High group shows the strongest negative correlation, with a rho value of -0.644 (p-value 1.01 x 10^-157^), followed by the Low group (rho value -0.624, p-value 2.75 x 10^-72^). The None group had the weakest correlation of the three, but this correlation was still strong and negative (rho value -0.615) and, like the High and Low groups, was highly significant (p-value 1.29 x 10^-41^). Regardless of the tRNA group for *Mycobacterium* phages, there appears to be strong selective pressure affecting codon bias, with codon bias decreasing in strength as GC3 decreases.

#### Correlation between EnC and GC3 by lifestyle according to host genus

Finally, comparing the correlation of EnC and GC3 in temperate and virulent phages can provide insight into the impact of a phage’s lifestyle on the strength of the selective pressure influencing codon bias. Nc plots showing the comparison between virulent and temperate phages per host genus are shown in **Figure 6**. As shown on the top row, *Escherichia* phages show a moderate positive correlation between EnC and GC3 for virulent phages that is highly significant (rho value 0.341, p-value 3.29 x 10^-91^), while the temperate phages show a weak negative correlation between these variables that is also significant (rho value -0.197, p-value 0.0052). A similar trend is seen in *Shigella* phages, where virulent phages have a moderate positive correlation between EnC and GC3 (rho value 0.302, p-value 4.44 x 10^-58^), but a weak negative correlation in temperate phages (rho value -0.190, p-value 4.88 x 10^-04^). In *Salmonella* phages, there was a very weak correlation between EnC and GC3 for both virulent and temperate phages (rho values 0.072 and 0.083 respectively), but only the correlation for virulent phages was significant (p-value 7.40 x 10^-05^). For Gram-negative bacteria in general, it seems that virulent phages have stronger relationships between EnC and GC3 than temperate phages, and that these relationships are positive. These findings indicate that as EnC increases (and codon bias decreases), the GC3 content also increases. Conversely, as GC3 content decreases, EnC decreases and codon bias increases, consistent with an AT-rich genome.

**Figure 6.**
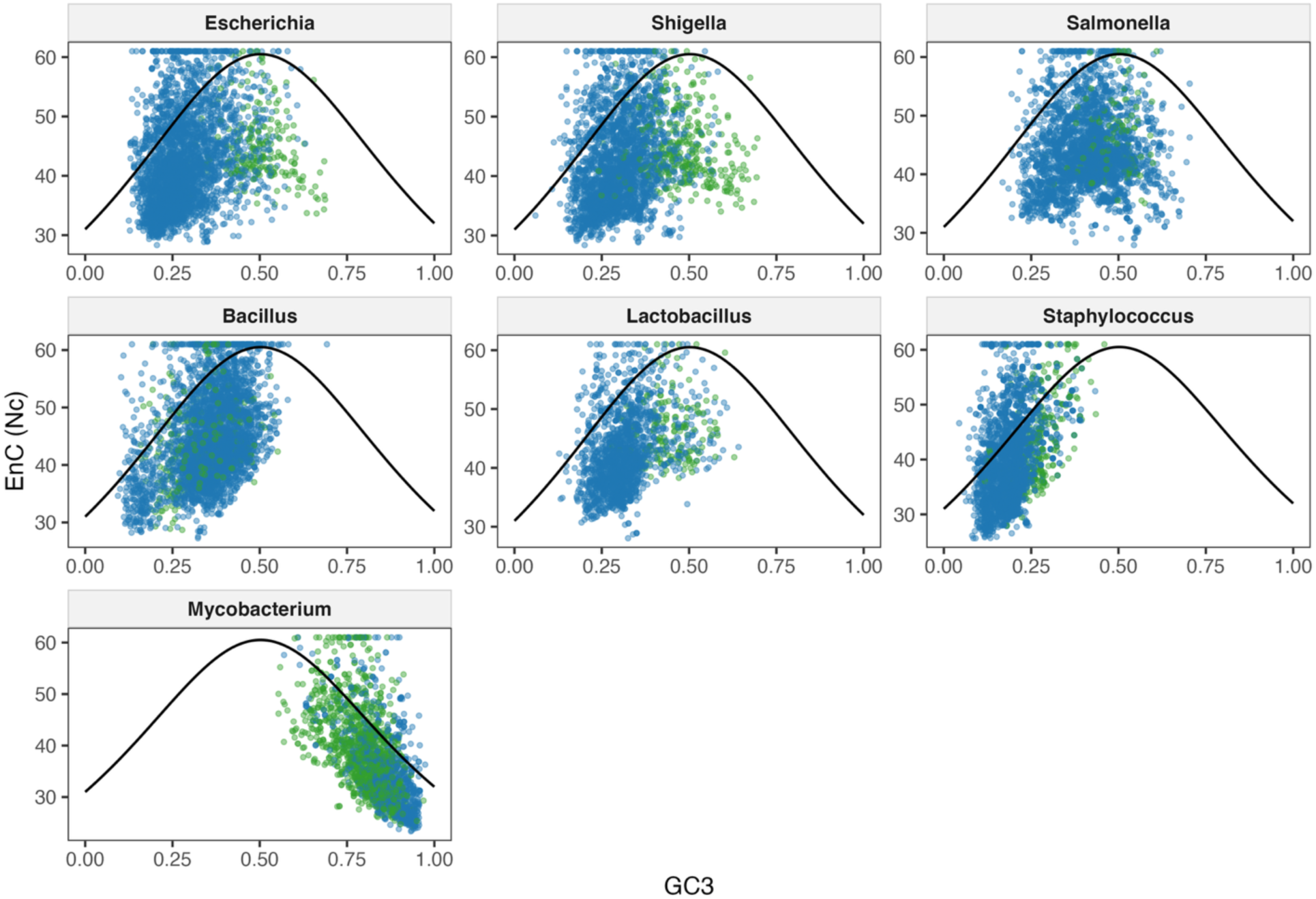
Nc plot of phages according to infection cycle. The Nc plots are faceted by host genus and depict the relationship between EnC and GC3 for each phage in each lifestyle group (Virulent or Temperate). The plot shows datapoints from all genes in all phages infecting that genus. Virulent phages are colored in blue, temperate phages are colored in green.

In *Lactobacillus* phages, a similar trend to the Gram-negative bacteria was found, with a moderate significant positive relationship between EnC and GC3 for virulent phages (rho value 0.367, p-value 3.11 x 10^-56^), and a very weak nonsignificant negative correlation for temperate phages (rho value -0.003). This contrasts with the *Bacillus* phages, which show nearly the same strength of relationship for both virulent and temperate phages (rho values 0.376 and 0.371, respectively), showing a moderate significant positive relationship between EnC and GC3 for both virulent and temperate phages (p-values 5.01 x 10^-117^ and 7.90 x 10^-09^, respectively). *Staphylococcus* phages were the only phages for which the temperate phages had a stronger relationship between EnC and GC3 than the virulent phages. Both virulent and temperate *Staphylococcus* phages showed positive moderate correlations between EnC and GC3 (rho values 0.398 and 0.495 respectively), both of which were highly significant (p-values 1.55 x 10^-118^ and 5.40 x 10^-16^ respectively). These results for these bacteria are shown in the middle row of **Figure 6**.

Perhaps unsurprisingly, the *Mycobacterium* phages (**Figure 6**, bottom row) show a strong negative correlation between EnC and GC3 for both virulent (rho value -0.628) and temperate (rho value - 0.586) phages. These relationships were also highly significant for both virulent and temperate phages (p-values 2.51 x 10^-128^ and 3.48 x 10^-114^ respectively). For Gram-positive phages, there seems to be less consistency in patterns of EnC and GC3 correlation between lifestyle groups, with some hosts showing stronger correlations with temperate phages, others with virulent phages, and still others with similar correlation strengths in both groups. For all but *Mycobacterium* phages, the significant correlations were positive, indicating that as codon bias increases, GC3 content decreases. Conversely, the *Mycobacterium* phages show trends of increasing codon bias as GC3 increases, likely due to the high GC content of the host and phage genomes.

### Relative Synonymous Codon Usage (RSCU) of Phages Compared to their Respective Hosts

Relative synonymous codon usage is a way of measuring the codon preference in a genome at single amino acid resolution (46). RSCU analyzes the frequency of the usage of each synonymous codon for a given amino acid and returns a value between 0, meaning that codon is never used, and up to however many synonymous codons there are for that amino acid. For example, the maximum value for a codon encoding leucine is 6, which would signify that only that codon is used for leucine, while the maximum value for one encoding proline is 4. Since methionine and tryptophan each only have one codon (CAT and TGG, respectively), they were not included in these analyses, as all RSCU values would be equal to either 1 (the codon is used) or 0 (the codon is absent). By comparing the RSCU of a phage with its host, codon bias differences can be seen at single-codon resolution. Because it is the relative usage, the usage pattern of amino acids in a gene does not impact the values. A heatmap depicting the mean RSCU values of each host genus and their phages is available in **Figure 7**, with Gram-negative bacteria and phages at the top and Gram-positive bacteria and phages at the bottom.

**Figure 7.**
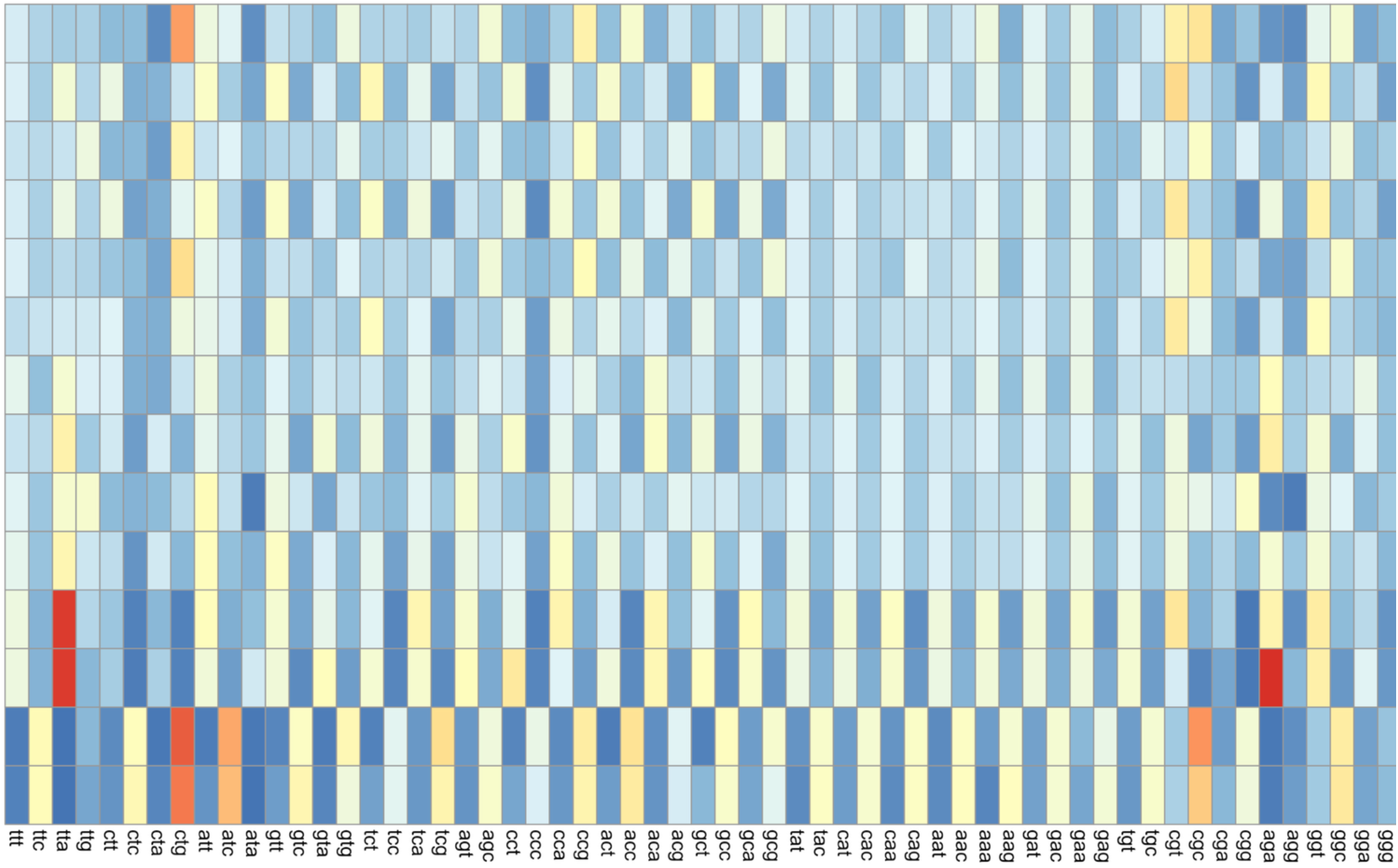
Heatmap of host and phage median RSCU for all codons. The median RSCU values for each host genus and their respective phages are plotted against each individual codon (excluding ATG and TGG). Lower values are in blue, while higher values are in red.

#### RSCU comparison of Gram negative hosts and their phages

Gram-negative bacteria in general had more significant differences in their codon usage compared to the phages that infect them. For *Escherichia* phages, 52 out of 61 coding codons had significantly different usage in the host compared to the phages (p-values range from 6.20 x 10^-05^ to 0.0304). Of these, the *Escherichia* phages collectively encode unique tRNAs that correspond with 21 of these significant codons, while only four of the unique tRNAs matched nonsignificant codons or singleton codons (specifically methionine). The *Shigella* phages had the highest number of significantly different codons when RSCU was compared to the host, with 57 codons showing significant differences (p-values range from 6.37 x 10^-05^ to 0.0455); only the codons GAA and GAG, both encoding glutamine, showed nonsignificant differences. The *Shigella* phages collectively encoded unique tRNAs corresponding to 23 of the significant codons, with only two tRNAs corresponding to nonsignificant or single codons. Thus, for both *Escherichia* and *Shigella* phages, tRNAs may compensate for these significant codon usage differences.

The *Salmonella* phages have fewer significant differences in their codon usage compared to their hosts, with only 39 codons showing significant differences in RSCU (p-values range from 1.54E-04 to 0.0320). Of these significantly different codons, 17 had unique corresponding tRNAs encoded by the *Salmonella* phages. An additional 13 unique nonsignificant or singleton-codon tRNAs are also encoded by these phages. In *Salmonella* phages, tRNAs may compensate for codon usage differences, but given the number of nonsignificant codons with corresponding tRNAs encoded, codon compensation may not be the only driving force behind tRNA acquisition.

#### RSCU comparison of Gram-positive hosts and their phages

The *Bacillus* phages had the highest number of significantly different codon RSCU values out of all of the Gram-positive hosts, with 51 out of 61 coding codons having significantly different usage in the phages compared to the host (p-values range from 1.71 x 10^-04^ to 0.0385). Collectively, the *Bacillus* phages encode unique tRNAs corresponding to 16 of these significantly different codons, with an additional eight unique tRNAs corresponding to nonsignificant or singleton codons (both methionine and tryptophan). In *Bacillus* phages, codon usage bias may be a driving force behind tRNA acquisition, in agreement with the EnC data.

With a similar number of codons showing significantly different RSCU values, *Staphylococcus* phages have significant differences for 46 codons (p-values range from 1.76 x 10^-04^ to 0.0325). The *Staphylococcus* phages encode only a small number of unique tRNAs, having only five unique tRNAs across all phages investigated. Two of these tRNAs have corresponding codons with significant differences in usage compared to the host, while three tRNAs correspond to nonsignificant or single codons. Because so few tRNAs are encoded in these phage genomes, it is difficult to determine if codon usage bias drives their acquisition.

*Lactobacillus* phages showed significant RSCU differences from their host for only 38 codons (p-values range from 0.00135 to 0.0325). Of these, nine have corresponding unique tRNAs encoded collectively by *Lactobacillus* phages, while eight nonsignificant or singleton codons have corresponding unique tRNAs. Like the *Salmonella* phages, there are similar numbers of tRNAs corresponding to significantly and nonsignificantly different codons, indicating that while codon usage bias may play a role in tRNA acquisition for these phages, it does not necessarily explain the presence of all tRNA genes.

Once again, *Mycobacterium* phages are the outlier, having the fewest number of codons showing significant RSCU differences between the phages and the host, with only 29 out of 61 coding codons showing significantly different RSCU values (p-values range from 8.35 x 10^-04^ to 0.0486). Of these 29 significantly different codons, 16 have unique corresponding tRNAs encoded by the *Mycobacterium* phages, while 21 of the unique tRNAs correspond with nonsignificant or singleton codons. The *Mycobacterium* phages also encode the highest number of unique tRNAs, with 37 different isodecoders encoded collectively. Since 16 of the 29 codons with significantly different RSCU values having corresponding tRNAs encoded by *Mycobacterium* phages, codon bias differences may drive the acquisition and/or retention of some of these tRNAs, contradicting previous studies (4,17,30).

### ΔRSCU Comparison of Phages According to tRNA Group

The ΔRSCU was calculated for each codon by subtracting the host mean RSCU for that codon from the phage mean RSCU for the same codon. These data were then used to quantify the differences between phage and host RSCU values according to how many tRNAs a phage encodes. The ΔRSCU was used instead of the RSCU to account for genome compositional differences of the hosts. Phages were again grouped into High (> 5 tRNAs), Low (1 – 5 tRNAs), and None (0) groups. Comparisons were made for High vs. Low, High vs. None, and Low vs. None groups, both within each host genus and globally. A High + Low combination group was also compared to the None group to examine the effects of having tRNAs in general on RSCU. A heatmap depicting the global ΔRSCU values for each codon according to phage tRNA group is shown in **Figure 8**.

**Figure 8.**
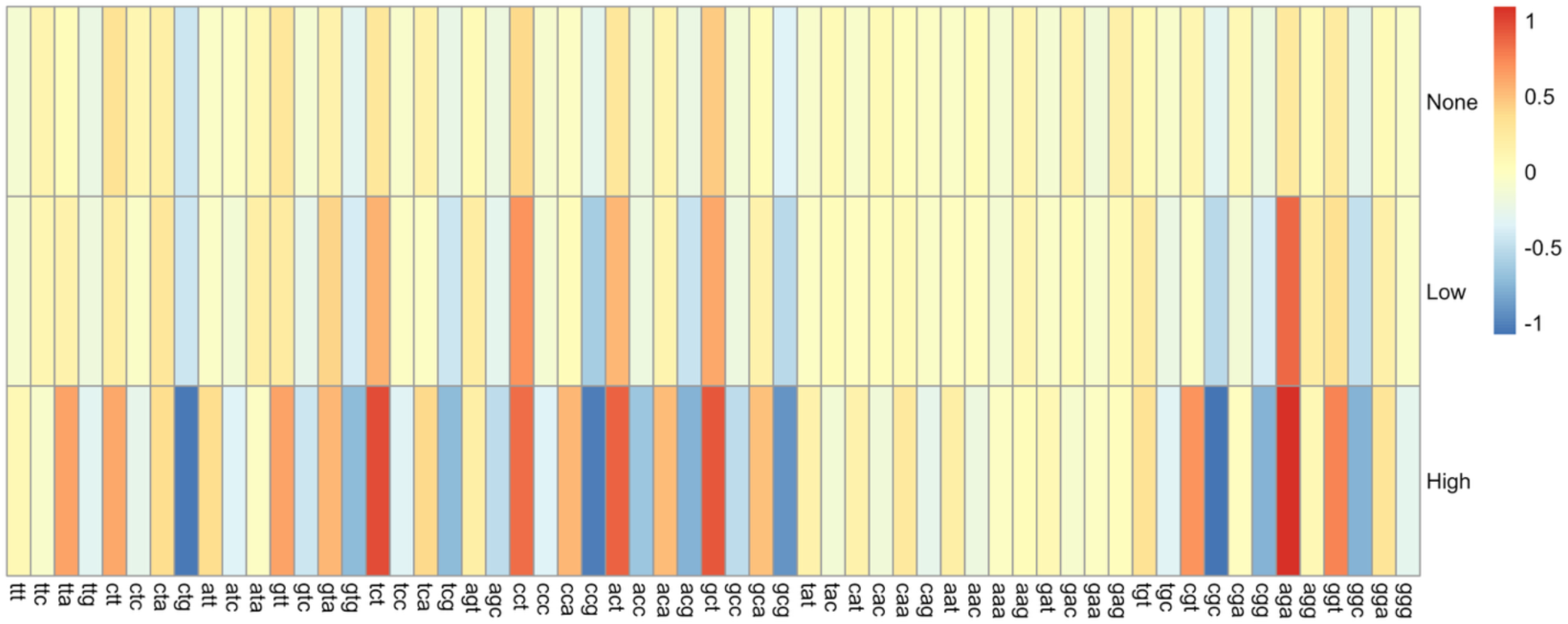
Heatmap of global ΔRSCU for all phages according to tRNA group. The median ΔRSCU values were calculated from the global pooled phage data for each tRNA group (High, Low, None). Low values in blue indicate less usage of that codon in the phage compared to the host. High values in red indicate more usage of that codon in the phage compared to the host.

#### ΔRSCU comparison of tRNA groups within each genus

The ΔRSCU of phages infecting a specific genus were compared according to the number of tRNAs encoded in each genome. As outlined above, tRNA groups were first compared individually, followed by a comparison of phages encoding tRNAs in general and those encoding none. For the *Escherichia* phages, 45 codons showed significantly different ΔRSCU values between phages that encode tRNAs and those that don’t (p-values range from 0.0057 to 0.0479). Of these 45 codons, 21 had corresponding unique tRNAs encoded collectively in the *Escherichia* phage genomes. Based on individual group comparisons, a total of 42 unique codons showed significant differences in ΔRSCU (p-values range from 3.96 x 10^-04^ to 0.0407). The High vs. None group comparison showed 37 codons with significant ΔRSCU differences between them, with 15 having corresponding tRNAs encoded collectively by the *Escherichia* phages. There were five codons with significantly different ΔRSCU values in the Low vs. None comparison, with two of these having corresponding tRNAs encoded by *Escherichia* phages. No codons were significantly different when the High vs. Low groups were compared. Overall, the largest ΔRSCU values were seen in the High group, indicating that higher numbers of tRNAs are associated with more deviation from the host RSCU values. Given the high number of tRNAs encoded by *Escherichia* phages, it is reasonable to conclude that these tRNAs compensate for codon bias differences.

The *Shigella* phages showed fewer significant differences in ΔRSCU overall, with 29 codons having significant differences when comparing phages with and without tRNAs (p-values range from 0.0139 to 0.0466). Of these 29 significantly different codons, only eight had corresponding tRNAs encoded collectively by the *Shigella* phages. When individual group comparisons were done, however, more codons show significant differences in ΔRSCU. A total of 43 unique codons showed significant differences in ΔRSCU when the groups were compared pairwise (p-values range from 4.38 x 10^-04^ to 0.0490). When the High vs. Low groups were compared, 29 codons showed significant differences in ΔRSCU values, with the High group consistently showing larger values and thus larger differences from the host. Of these 29 codons, eight had corresponding tRNAs encoded by the *Shigella* phages. When the High vs. None groups were compared, 36 codons showed significant differences in ΔRSCU, with the High group again showing larger values. Of these, 11 had corresponding tRNAs encoded by the *Shigella* phages. Conversely, when the Low vs. None groups are compared, only two codons showed significant differences, neither of which had a corresponding tRNA found in the phage genomes. For *Escherichia* phages, having tRNAs in general seems to be associated with greater differences in codon usage; however, the number of tRNAs is more important in *Shigella* phages, with phages encoding high numbers of tRNAs consistently showing larger differences in RSCU compared to the host than phages encoding few or no tRNAs.

While the *Salmonella* phages showed no codons with significant ΔRSCU differences when phages with and without tRNAs were compared, there were 51 significantly different codons when the individual pairwise comparisons of the groups were performed (p-values range from 8.59 x 10^-04^ to 0.0488). The largest number of codons with significantly different ΔRSCU values was seen in the High vs. Low group comparison, with 44 codons having significant differences. Of these, 20 have corresponding tRNAs collectively encoded in the *Salmonella* phage genomes. These differences may explain why no significance was seen when comparing phages with and without tRNAs, as most of the significant differences in codons occur when two groups with tRNAs are compared. When the High vs. None groups are compared, 25 codons show significant ΔRSCU differences, with eight having corresponding tRNAs in the *Salmonella* phages. Lastly, 14 codons show significant differences in ΔRSCU when the Low vs. None groups are compared, and 10 of these have corresponding tRNAs encoded by the *Salmonella* phages. Unlike *Escherichia* and *Shigella* phages, where there is a clear pattern of the High group having consistently larger ΔRSCU values, there is more variation in the *Salmonella* phage data. For some codons, the High group shows the largest values. For others, it is the Low group or even the None group. This makes it difficult to draw conclusions on the impact of having tRNAs and/or the number of tRNAs in the genome on the magnitude of ΔRSCU.

In terms of Gram-positive hosts, the *Bacillus* phages showed no significant differences in ΔRSCU for all codons when phages with and without tRNAs were compared. When individual pairwise comparisons were performed, however, 27 codons showed significant differences in ΔRSCU (p-values range from 0.0015 to 0.0459). Specifically, there were 22 codons with significantly different ΔRSCU values when the High vs. Low groups were compared, nine of which have corresponding tRNAs encoded collectively by the *Bacillus* phages. The High vs. None group comparison showed 15 codons with significantly different ΔRSCU values, and of these, seven have corresponding tRNAs in the *Bacillus* phage genomes. Lastly, eight codons showed significant differences in ΔRSCU when the Low vs. None groups were compared, one of which has a corresponding tRNA in the *Bacillus* phage genomes. Overall, more codons showed the largest ΔRSCU values in the High group. However, like the *Salmonella* phages, many codons also showed the highest ΔRSCU values for the Low or None group. This makes it challenging to determine the relationship between number of phage-encoded tRNAs and the differences in phage vs. host RSCU.

The *Lactobacillus* phages also showed no significant differences in ΔRSCU for any of the codons when comparing phages with tRNAs to those without, but 18 codons showed significant differences in ΔRSCU when the pairwise comparisons of the individual groups were performed (p-values range from 0.0139 to 0.0463). When comparing the High vs. Low groups, 10 codons showed significant differences in ΔRSCU, with one codon having a corresponding tRNA encoded by the *Lactobacillus* phages. There were 17 codons with significantly different ΔRSCU between the High vs. None groups, with three having corresponding tRNAs encoded by the *Lactobacillus* phages. Conversely, none of the codons in the Low vs. None group comparison showed any significant differences in ΔRSCU. Many of the codons show the largest ΔRSCU values in the High group, indicating phages with high numbers of tRNAs have larger differences from the host RSCU than those with low or no tRNAs.

Likewise, the *Mycobacterium* phages showed no significant differences in ΔRSCU for any of the codons when comparing phages with tRNAs to those without tRNAs, with 18 codons showing significant differences when performing pairwise comparisons on the tRNA groups (p-values range from 0.0021 to 0.0443). Unlike the *Lactobacillus* phages, nearly all codons with significant differences in ΔRSCU values had corresponding tRNAs in the *Mycobacterium* phage genomes. When comparing the High vs. Low groups, there were 12 codons showing significant differences in ΔRSCU, 11 of which have corresponding tRNAs in the *Mycobacterium* phage genomes. Similarly, the High vs. None groups have nine codons showing significant differences in ΔRSCU between them, seven of which have corresponding tRNAs in the phage genomes. There is one codon significantly different in ΔRSCU between the Low vs. None groups, and this codon has a corresponding tRNA in the phage genomes. When looking at the ΔRSCU data for the groups, no clear pattern emerges as to which tRNA group differs in RSCU the most from the host. It is likely though that these tRNAs compensate for codon usage differences, as the High group typically showed the largest ΔRSCU values specifically for the codons showing significant differences.

The *Staphylococcus* phages were unique in that none of the 23 phages investigated had more than 4 tRNAs, so the only pairwise comparison that could be performed was between the Low vs. None groups, which also by default was the comparison between phages with and without tRNAs. When these groups were compared, 36 codons showed significant differences in ΔRSCU (p-values range from 0.0172 to 0.0455), with no clear pattern as to which group showed larger ΔRSCU values. Additionally, only two of these codons have corresponding tRNAs encoded in the *Staphylococcus* phage genomes. In this case, it is difficult to draw conclusions on whether tRNAs are compensating for codon usage differences.

#### ΔRSCU comparison of tRNA groups globally across genera

When pooling all phage ΔRSCU data, 38 codons showed significantly different ΔRSCU values in phages with tRNAs compared to those without (p-values range from 3.92 x 10^-06^ to 0.0434). Phages with tRNAs tended to have larger ΔRSCU values, and thus larger RSCU differences compared to their respective hosts. When the individual tRNA groups were compared against each other, a total of 54 codons showed significant differences in ΔRSCU. The High vs. None comparison yielded the highest number of significantly different codons, with 52 codons showing significant differences in ΔRSCU. The High vs. Low group comparison found 47 codons with significant ΔRSCU differences. There were 20 significantly different codons in the Low vs. None comparison. Overall, the High group tended to have the largest ΔRSCU values, indicating that not only the presence, but also the number of tRNAs is associated with larger differences in codon bias in phages compared to their hosts on a global scale across all genera investigated.

### ΔRSCU Comparison According to Lifestyle

In addition to the effect of tRNA numbers on the differences in phage vs. host codon usage bias, the phage lifestyle was also investigated to determine its impact on ΔRSCU values. Virulent and temperate phages were compared within each host genus and globally across host genera, as before. A heatmap depicting the global ΔRSCU values for each codon according to phage lifestyle is shown in **Figure 9**.

**Figure 9.**
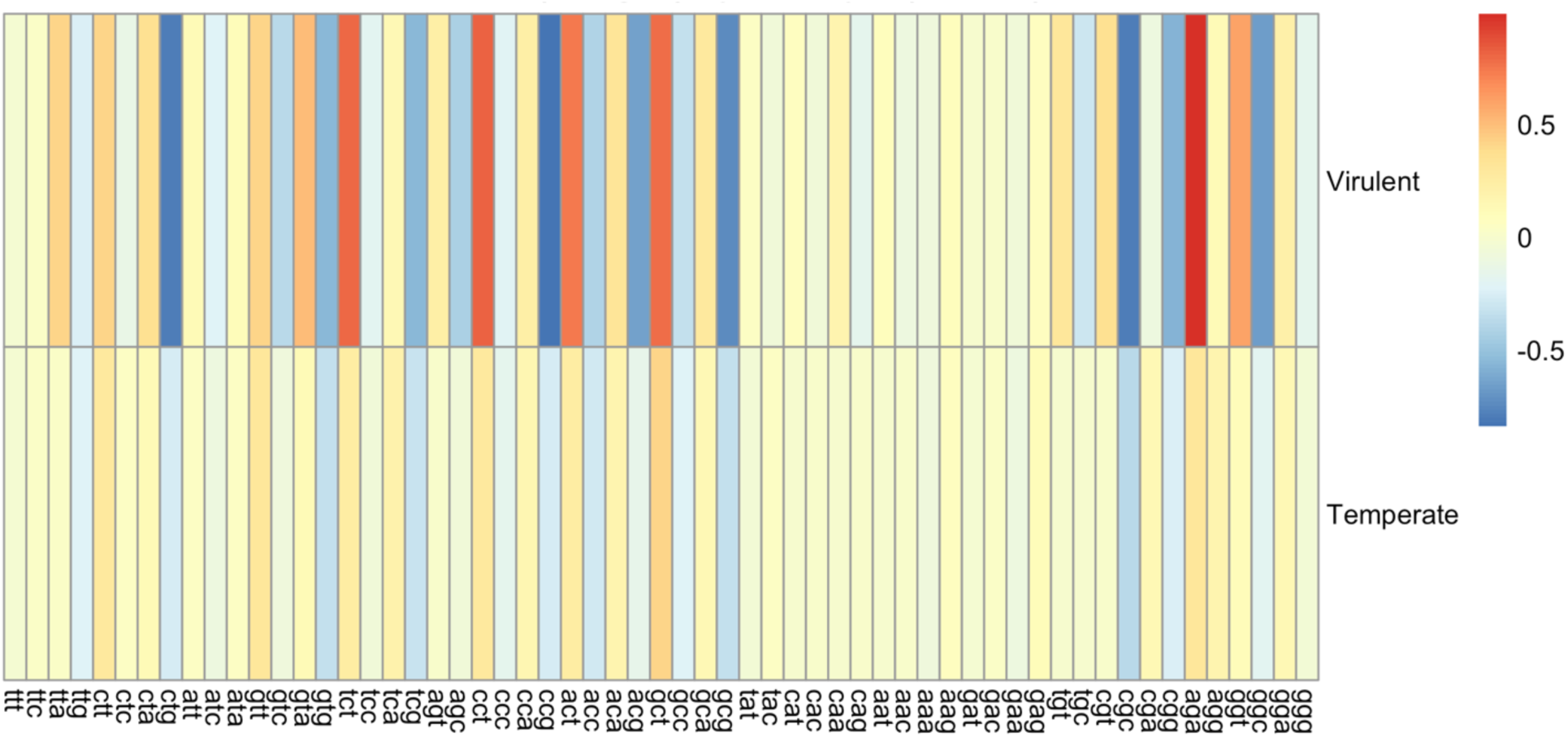
Heatmap of global ΔRSCU for all phages according to lifestyle. The median ΔRSCU values were calculated from the global pooled phage data for each lifestyle (Virulent or Temperate). Low values in blue indicate less usage of that codon in the phage compared to the host, High values are red and indicate more usage of that codon in the phage compared to the host.

#### ΔRSCU comparison of lifestyle within a host genus

Temperate and virulent *Escherichia* phages had 28 codons with significantly different ΔRSCU (p-values range from 0.0077 to 0.0491). Of these, nine had corresponding tRNAs collectively encoded by the *Escherichia* phages. Nearly all codons showed higher absolute ΔRSCU values in virulent phages, and the few codons without higher ΔRSCU did not show significant differences between lifestyles. Since there was only one tRNA-encoding temperate *Escherichia* phage included in this analysis (phage P1), it is possible that the phage-encoded tRNAs are used to compensate for codon usage differences between the host and virulent phages. Conversely, the *Salmonella* phages showed no significant differences in ΔRSCU for any of the codons when temperate and virulent phages were compared, suggesting this has little effect on the codon usage differences between a phage and its host. As mentioned earlier, it should be noted that only two temperate phages infecting *Salmonella* met our criteria for analysis, so as with *Escherichia*, this may not necessarily be reflective of all *Salmonella* phages.

The *Shigella* phages had the largest number of significantly different codons, with 45 out of 61 codons having significant differences in ΔRSCU (p-values range from 4.92 x 10^-04^ to 0.0460). Of these codons, 16 have corresponding tRNAs collectively encoded in the phage genomes. For the majority of significantly different codons, the ΔRSCU were larger in virulent phages. Like temperate *Escherichia* phages, temperate *Shigella* phages showed only a few codons with larger ΔRSCU values, three of which were significantly different. Only the temperate phage Sf6 encoded tRNAs; otherwise, tRNAs were almost exclusively found encoded in the virulent phage genomes. Both tRNAs encoded by Sf6 recognize codons that are significantly different between the host and phage, but neither of these codons had larger ΔRSCU values for temperate phages in general. However, many of the virulent phages encoded large numbers of tRNAs, with 10 phages belonging to the family *Ounavirinae.* These phages encoded 25-27 tRNAs, which recognize a nearly full repertoire of amino acids, with the exception of tryptophan. This notable difference in tRNA number between temperate and virulent phage genomes suggests that phage-encoded tRNAs compensate for codon usage bias differences between the host and virulent phages specifically.

The *Bacillus* phages had only one codon with significantly different ΔRSCU values when comparing temperate with virulent phages (p-value 0.0295). These phages collectively encode a tRNA corresponding to the codon CTA (Leu), which was the singular significantly different codon between groups. Similarly, *Lactobacillus* phages show no significant differences in ΔRSCU when comparing temperate and virulent phages. For these hosts, lifestyle does not seem to impact how different a phage’s codon usage is from its host. Results mentioned above also showed no significant differences in ΔRSCU when comparing phages with or without tRNAs, but nearly all codons showed significant differences between phages and their hosts. Combined with these lifestyle data, it appears that codon usage bias of all *Bacillus* phages are significantly different from their host bacteria. This difference is seen across all phages of these hosts, rather than in specific groups of phages. The same pattern is seen in the *Lactobacillus* phages, suggesting the codon usage bias differences are common in all phages regardless of lifestyle or whether a phage encodes tRNAs.

The *Staphylococcus* phages showed patterns closer to the Gram-negative hosts *Escherichia* and *Shigella*, with 37 codons having significant differences in ΔRSCU between the virulent and temperate phages (p-values range from 0.0053 to 0.0390). Of these, two codons have corresponding tRNAs encoded collectively by the *Staphylococcus* phages – AAC (Asn; p-value 0.0053) and GAC (Asp; p-value 0.0081). These are the same tRNAs that match the codons with significant ΔRSCU differences in the tRNA group comparison, plus the comparison of RSCU between host and phage. Unlike the phages infecting Gram-negative hosts, there was no consistent pattern regarding which group had larger absolute ΔRSCU values, with approximately half of all codons having larger ΔRSCU values for temperate phages, and others having larger ΔRSCU values for virulent phages. This pattern was also seen when looking at significant codons only. While lifestyle does impact the codon usage of these phages significantly, the magnitude of that difference does not seem to be impacted by the lifestyle of that phage.

Similar to the *Staphylococcus* phages, the *Mycobacterium* phages had many codons with significant differences in ΔRSCU when comparing virulent and temperate lifestyles. Of the 29 codons with significant ΔRSCU differences (p-values range from 6.56 x 10^-04^ to 0.0342), 19 have corresponding tRNAs encoded in the *Mycobacterium* phage genomes. Also like the *Staphylococcus* phages, approximately half of all codons have higher absolute ΔRSCU values in the temperate phages, while the others have the highest absolute values in the virulent phages. When looking specifically at significant codons, this same pattern is seen. As mentioned above, *Mycobacterium* phages with or without tRNAs also showed no differences in codon usage when comparing between the host and the phages. These results indicate that codon bias differences between *Mycobacterium* phages and their host are not tRNA-dependent but codon usage patterns may depend on lifestyle. Since both temperate and virulent *Mycobacterium* phages can encode large repertoires of tRNAs, this could explain why some of the significant codons show larger ΔRSCU values in temperate phages, while others show larger values in virulent phages.

#### ΔRSCU comparison of lifestyle globally across genera

When comparing the ΔRSCU values of temperate and virulent phages on a global scale across genera, virulent phages consistently had larger absolute ΔRSCU values compared to temperate phages and thus showed greater differences from their host codon usages. A total of 37 codons showed significant differences in ΔRSCU between virulent and temperate phages (p-values range from 4.90 x 10^-11^ to 0.0321), with only two of the significant codons having larger ΔRSCU values compared to the virulent phages – CGA (Arg: p-value 6.67 x 10^-04^) and CTC (Leu; p-value 3.64 x 10^-05^). In total, only 11 codons had larger ΔRSCU values for temperate phages, indicating that virulent phages in general have greater disparity in codon usage compared to their hosts than temperate phages. These results overall were also highly significant.

### PERMANOVA on ΔRSCU Variables

A PERMANOVA was run on the ΔRSCU data to determine if the ΔRSCU profiles differ by lifestyle and/or tRNA group. The R2 value measures the total variance in ΔRSCU profiles based on the grouping factors, while the F statistic represents the ratio of between-group variation to within-group variation and is used to determine significance. The model used the Δ distance, calculated by subtracting the host mean RSCU values from the phage mean RSCU values per codon; lifestyle; and tRNA group for the within-genus comparisons. When the PERMANOVA was applied globally, host genus was also added to the model.

#### PERMANOVA in Gram negative hosts

Phages infecting the Gram-negative hosts were examined first. For the *Escherichia* phages, the PERMANOVA showed significant differences in ΔRSCU patterns, indicating that these differences are significantly affected by both tRNA group (R2 = 0.265, F = 5.518, p-value = 0.0010) and lifestyle (R2 = 0.141, F = 5.863, p-value = 0.0024). Together, these factors explained ∼40% of the variance seen. Similar results were seen with the *Shigella* phages, with both the tRNA group (R2 = 0.177, F = 3.931, p-value = 0.004) and lifestyle (R2 = 0.082, F = 3.643, p-value = 0.016) having significant effects on the ΔRSCU profiles of the *Shigella* phages, together explaining ∼26% of variances. The *Salmonella* phages showed similar results, with the PERMANOVA showing significant effects of both tRNA group (R2 = 0.486, F = 10.559, p-value = 0.0002) and lifestyle (R2 = 0.082, F = 3.546, p-value = 0.0128) on the ΔRSCU profiles, together explaining ∼57% of variance seen in ΔRSCU patterns. For all genera, the tests for homogeneity were nonsignificant for both tRNA and lifestyle, indicating that the differences in ΔRSCU patterns were due to compositional differences in ΔRSCU profiles rather than unequal variance of the groups.

#### PERMANOVA in Gram positive hosts

Phages infecting Gram-positive bacteria were analyzed next. Like the Gram-negative genera, effects of phage tRNA group and lifestyle were significant for Gram-positive genera. Specifically, the PERMANOVA on the *Bacillus* phages showed a significant effect of phage tRNA group (R2 = 0.212, F = 3.076, p-value = 0.002) and lifestyle (R2 = 0.103, F = 3.001, p-value = 0.017) on the ΔRSCU profiles of the phages, together explaining ∼31% of variance seen. However, the tests for homogeneity revealed significant differences among the tRNA groups (F = 3.741, p-value = 0.0354), indicating the results of the PERMANOVA may be influenced by variance within the tRNA groups, rather than true compositional differences in ΔRSCU across all variables.

The PERMANOVA on *Lactobacillus* phages showed significance for tRNA group only (R2 = 0.274, F = 3.124, p-value = 0.0156), explaining ∼27% of variance in ΔRSCU patterns. For these phages, lifestyle did not show significance. Conversely, as with all other groups, the PERMANOVA on *Mycobacterium* phages showed significance for both the tRNA group (R2 = 0.167, F = 2.374, p-value = 0.0046) and lifestyle (R2 = 0.183, F = 5.216, p-value = 0.0002), explaining ∼35% of variance in ΔRSCU profiles among phages. Additionally, for both *Lactobacillus* and *Mycobacterium*, the tests for homogeneity were nonsignificant for both tRNA group and lifestyle, indicating that the differences in ΔRSCU patterns were due to true compositional differences rather than variance within the groups.

In the *Staphylococcus* phages, lifestyle and tRNA group were perfectly confounded, with all temperate phages having no tRNAs and all virulent phages belonging to the Low tRNA group. Because of this, each factor (lifestyle and tRNA group) was tested individually. When tRNA group was used as the predictor, the PERMANOVA showed a significant effect on ΔRSCU patterns (R2 = 0.276, F = 7.996, p-value = 0.0012). Likewise, when lifestyle alone was used as the predictor, there was a significant impact of lifestyle on ΔRSCU patterns (R2 = 0.276, F = 7.996, p-value = 0.0012). The tests for homogeneity showed no significance, indicating these ΔRSCU pattern differences were due to true compositional differences and not due to variance within the groups.

#### Global PERMANOVA

Once these groups had been analyzed individually, all phage data were pooled together for a global PERMANOVA using the predictors of tRNA group, lifestyle, and host genus to determine whether these factors impact ΔRSCU profiles. The results indicate that both tRNA group (R2 = 0.065, F = 9.681, p-value = 0.0002) and lifestyle (R2 = 0.013, F = 3.952, p-value = 0.0040) have a significant effect on ΔRSCU patterns, together contributing to ∼8% of total variance. When the tests for homogeneity on the lifestyle and tRNA groups were performed, lifestyle showed high significance while tRNA showed nonsignificant results. This indicates that the global effect of lifestyle is due to variance within the groups rather than true compositional differences in ΔRSCU patterns between lifestyles. However, the variance seen from tRNA groups is due to true differences in ΔRSCU patterns between groups. Lastly, the host genus predictor did not show any significance in the PERMANOVA, indicating the host genus of a phage does not affect the ΔRSCU patterns. The results of the global PERMANOVA are visualized in the PCA plot in **Figure 10**.

**Figure 10.**
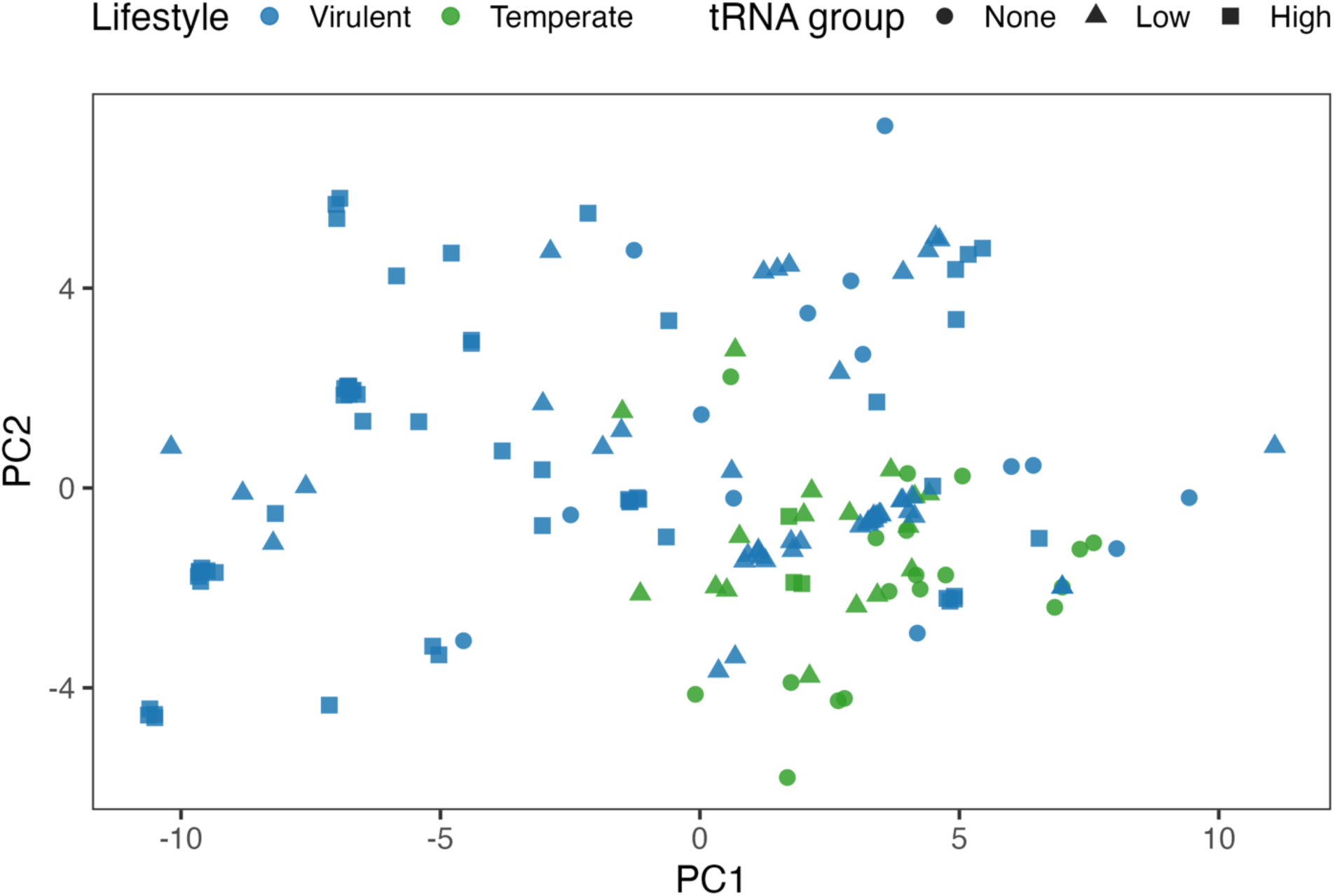
PCA plot of ΔRSCU vectors. Phages are colored by lifestyle (Virulent in blue, Temperate in green) and shaped according to tRNA group (High is a filled square, Low is a filled triangle, and None is a filled circle).

### tRNA Adaptation Index (tAI) of Phages Compared to their Respective Hosts

The tRNA adaptation index is a computational index estimating translational efficiency based on how well an organism’s codon usage is adapted to its tRNA pool (59–62). Values range from 0 to 1, with values closer to 0 showing lower levels of codon adaptation, and values closer to 1 showing higher levels of codon adaptation which is associated with increased translational efficiency. The tAI was calculated per gene for all phages and all hosts using each organism’s respective RSCU and the tRNA pool encoded by the host. Calculating tAI using RSCU values does not require prior information on gene expression levels or even functional annotation, which is an important feature when studying phages, as many genes are hypothetical. The resulting data estimates the translational efficiency of each gene in each organism, and these data were then used to determine the significance of the difference in phage median tAI compared to that of their hosts. The distributions of tAI values for each host genus are shown in **Figure 11**, and a statistical summary of the tAI data can be found in **Table 4**.

**Figure 11.**
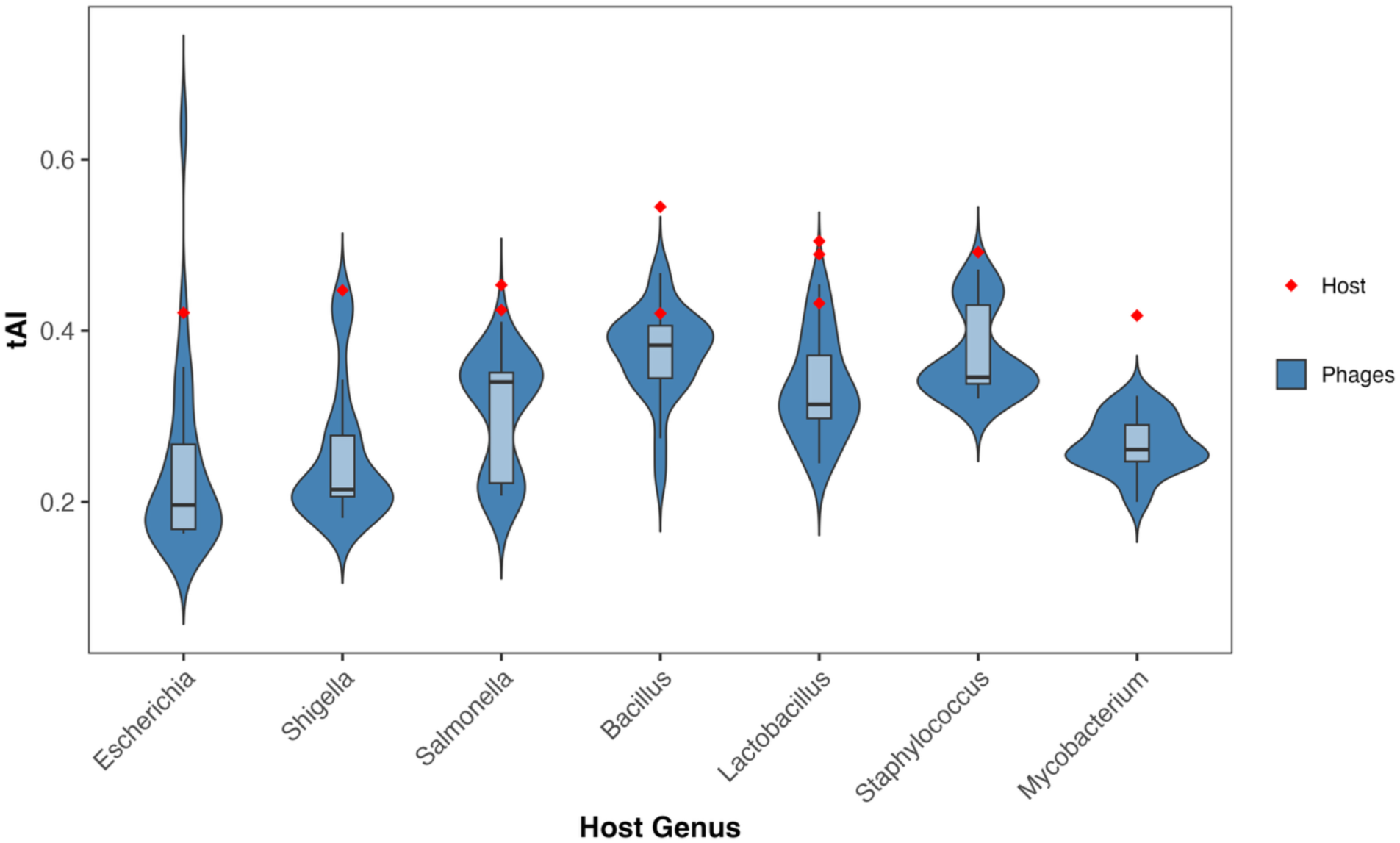
Distribution of median tAI values in phages according to host genus. The violin plot depicts the median tAI of each phage as a blue distribution, with the median host tAI as a red dot. For host genera where multiple species or serovars were used, multiple red dots appear.

**Table 4.** Summary of median tAI values in phages and hosts.

| Host Genus | N host | N phage | Median host | Median phage | Std dev tAI |
| --- | --- | --- | --- | --- | --- |
| *Escherichia* | 1 | 24 | 0.4210 | 0.1962 | 0.1119 |
| *Shigella* | 1 | 24 | 0.4473 | 0.2144 | 0.0789 |
| *Salmonella* | 2 | 22 | 0.4388 | 0.3401 | 0.0668 |
| *Bacillus* | 2 | 23 | 0.4825 | 0.3831 | 0.0528 |
| *Lactobacillus* | 3 | 17 | 0.4893 | 0.3138 | 0.0594 |
| *Staphylococcus* | 1 | 23 | 0.4918 | 0.3457 | 0.0509 |
| *Mycobacterium* | 1 | 21 | 0.4178 | 0.2611 | 0.0326 |

For all genera, both Gram-negative and Gram-positive, the differences in phage tAI values compared to those of their hosts were significant. Beginning with the phages infecting the Gram-negative genus *Escherichia*, the median tAI of the host (0.421) was significantly higher than the median tAI of the phages (0.196, p-value 1.47 x 10^-05^). *Escherichia* phages showed the smallest median tAI values, indicating these phages have the lowest level of adaptation to the host tRNA pool out of all host genera. The *Shigella* phages showed the highest significance between the host median tAI (0.447) and the phage median tAI (0.214, p-value 8.33 x 10^-07^) and the largest ΔtAI value (−0.233). These results show a large difference in the level of adaptation between phage and host, with the host being significantly more adapted to its own tRNA pool. Out of all host genera, *Shigella* phages showed the largest difference in adaptation level compared to their host. The *Salmonella* phages also showed highly significant differences between the host median tAI (0.439) and the phage median tAI (0.340, p-value 1.67 x 10^-06^). However, *Salmonella* phages also had the smallest ΔtAI values (median = -0.092). This is consistent with data from other indices, which showed more similarities in codon usage between the *Salmonella* and their phages compared to many of the other genus-phage pairs in this study.

In terms of Gram-positive genera, the differences in phage median tAI and their respective host median tAI were also highly significant. The *Bacillus* phages had the highest median tAI (0.383) of all the genera investigated, which is is significantly different from the median tAI of the host (0.483, p-value 4.17 x 10^-06^), resulting in a median ΔtAI of -0.136. The median tAI for the *Lactobacillus* was similar to that of *Bacillus* (0.489), but the median tAI of the *Lactobacillus* phages was lower (0.314) and the difference was also significant (p-value 1.47 x 10^-05^). The median ΔtAI of the *Lactobacillus* phages was -0.121. Out of all of the host genera, *Staphylococcus* showed the highest level of adaptation to its own pool (median tAI = 0.492). The median tAI for the *Staphylococcus* phages was significantly different from the host (0.346, p-value 2.88 x 10^-05^), with a resulting median ΔtAI of -0.146. *Mycobacterium* showed the lowest adaptation to its own tRNA pool (median tAI = 0.418), with the phage median tAI values being significantly lower (0.261, p-value 2.23 x 10^-06^). The median ΔtAI of the *Mycobacterium* phages was -0.157.

These data altogether indicate that the phage codon usage biases across all genera are significantly less adapted to their host tRNA pool than their host bacteria are. Lower levels of adaptation to the host pool may indicate a potential benefit to phages encoding tRNAs: a phage that encodes its own tRNAs to add to the host pool can boost translational efficiency. The *Shigella* phage Sf14 has been shown to have increased tAI when using only the phage tRNA pool (16), and it is known that the *Escherichia* phage T4 encodes tRNAs to increase its translational efficiency during infection (63). This phenomenon may also apply to other phages analyzed here.

### ΔtAI Comparison within a Host Genus

Because tAI values can vary greatly within genomes, the tAI values for each gene in the phage genomes were considered in the analysis. Unlike the previous indices used––which use a median value either per phage, or per codon per phage for calculations––the most accurate way to assess tAI is using the full data from the calculations rather than a median. Because of this, a mixed model was used to investigate the impacts of the number of tRNAs in a phage genome and/or a phage’s lifestyle on how different its tAI values are compared to its host. As before, the ΔtAI values were calculated by subtracting the tAI values of the respective host from that of the phage. The mixed model used the predictors of tRNA group and lifestyle to determine whether one or both factors systematically shift ΔtAI values, with the random effect set to the phage genome to account for multiple genes coming from the same genome. The baseline values for the mixed model for each predictor were “None” for the tRNA group and “Temperate” for the lifestyle. The intercept for each genus was calculated from the ΔtAI values of temperate phages and those with no tRNAs to determine if the baseline values are significantly different from 0. Logit-scaled values were then back-transformed into ΔtAI values for ease of interpretation. Significant negative values represent lower adaptation of the phage to the host’s tRNA pool versus the bacteria; significant positive values represent higher adaptation. The distribution of ΔtAI values for phages are faceted by host genus for both tRNA group and lifestyle comparisons in **Figure 12**.

**Figure 12.**
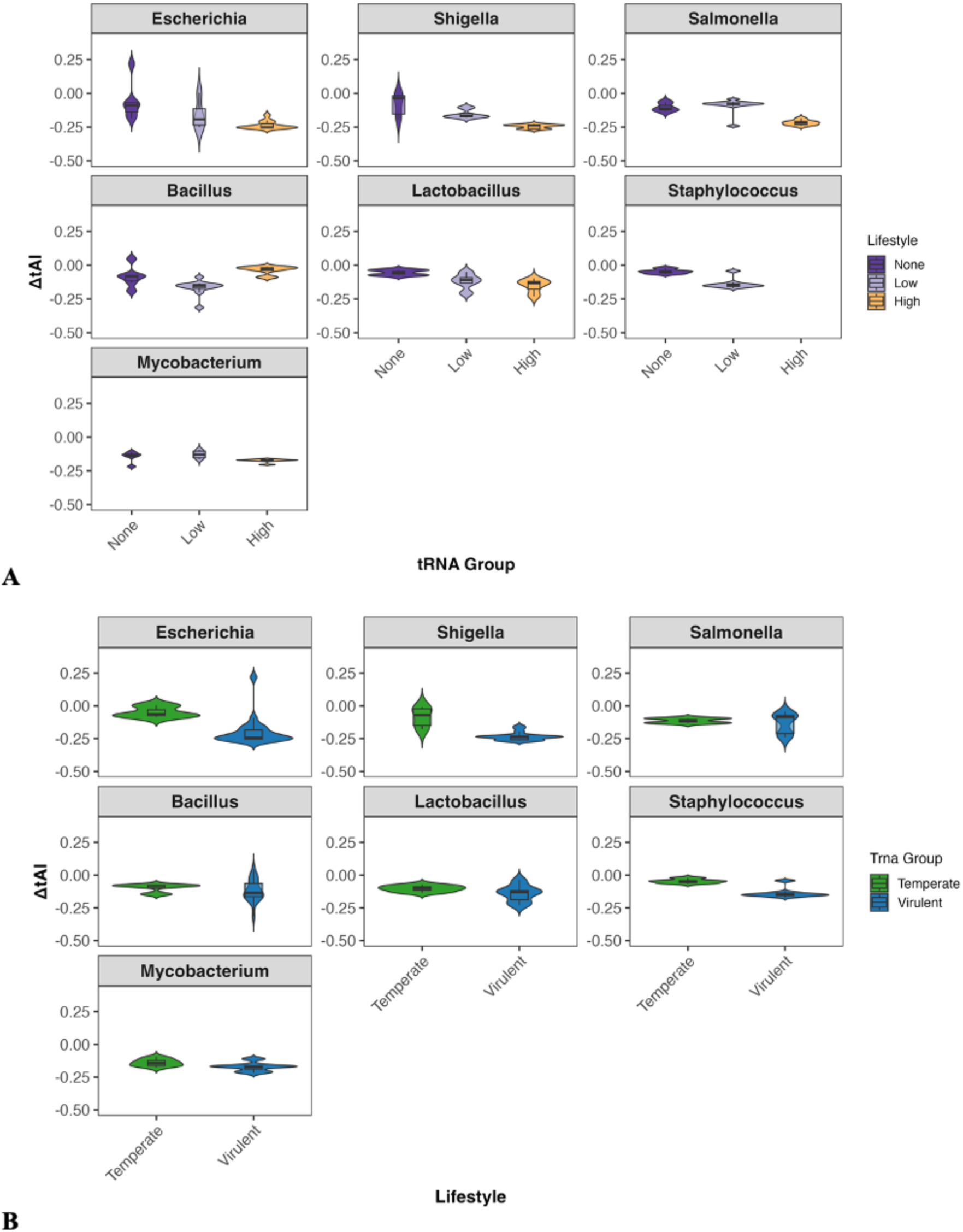
Distribution of ΔtAI in phages by bacterial genus. A) A violin plot faceted by host genus depicting the distributions of ΔtAI values in the phages belonging to each of the three tRNA groups (High, Low, None). High is colored in yellow, Low in light purple, and None in dark purple. B) A violin plot faceted by host genus depicting the distributions of ΔtAI values in the phages belonging to each lifestyle (Virulent or Temperate). Virulent are colored in blue, temperate in green

#### ΔtAI comparison in Gram-negative hosts

In *Escherichia* phages, the intercept was significant, with a logit-scaled mean of -0.363 (p-value 0.0063), indicating the temperate phages and those with no tRNAs have ΔtAI that are significantly less adapted to the host tRNA pool than the bacteria. When investigating the impact of tRNA group on how different a phage’s tAI values are compared to its host, the Low and High group were each individually compared to the baseline None group. When the Low group was compared to the None baseline, phages in the Low group had smaller ΔtAI values compared to the None group, but these difference in ΔtAI between phages were not significant. Conversely, the High group showed a significant difference in ΔtAI compared to the None group. The High group had a mean ΔtAI of -0.180, which was significantly larger than that of the None group (−0.094, p-value 0.0026). This indicates that *Escherichia* phages with high numbers of tRNAs are significantly less adapted to the host tRNA pool than those with none. When lifestyle was examined, virulent phages also showed a significantly larger mean ΔtAI (−0.196) compared to the temperate phages (−0.076, p-value 1.28 x 10^-04^), indicating that––like phages with high numbers of tRNAs––virulent phages are significantly less adapted to the host tRNA pool than temperate phages.

The *Shigella* phages showed significance for all groups compared. The logit-scaled intercept of the temperate and no tRNA phages was -0.372, significantly different from 0 (p-value 3.69 x 10^-05^). This indicates the phage baseline group is significantly less adapted to the host tRNA pool than the bacteria. In the tRNA comparisons, the mean ΔtAI of both the Low group (−0.184) and the High group (−0.196) were significantly different from that of the None group (−0.102, p-values 7.89 x 10^-06^ and 3.01 x 10^-09^, respectively). These data indicate that having tRNAs in general results in significantly lower adaptation to the host tRNA pool. Likewise, there was a highly significant difference in the mean ΔtAI for virulent phages (−0.205) compared to temperate phages (−0.116, p-value 7.85 x 10^-05^), indicating that like the *Escherichia* phages, virulent phages are significantly less adapted to the host tRNA pool than temperate phages.

In the *Salmonella* phages, the logit-scaled intercept of temperate and no tRNA phages was -0.782, significantly different from 0 (p-value 3.60 x 10^-06^). Again, this indicates the baseline group is significantly less adapted to the host tRNA pool than the host. When comparing the tRNA groups, only the High group had a mean ΔtAI significantly different from the None group (−0.211 vs. - 0.109, p-value 9.72 x 10^-04^). This indicates that while low numbers of tRNAs do not result in significantly different adaptation to the host tRNA pool than those with no tRNAs, high numbers of tRNAs result in significantly lower adaptation. Interestingly, the lifestyle of *Salmonella* phages does not seem to significantly affect the ΔtAI of a phage, though the temperate group showed slightly lower levels of adaptation to the host tRNA pool, with a mean ΔtAI of -0.158, compared to a mean ΔtAI of -0.129 for the virulent phages.

#### ΔtAI comparison in Gram-positive hosts

The *Bacillus* phages had a logit-scaled intercept of -0.307 for the temperate and no tRNA phage group, indicating this baseline group is significantly less adapted to the host tRNA pool than the host (p-value 0.0130). Unlike the Gram-negative hosts, the High tRNA group showed only a non-significantly smaller ΔtAI from the None group, indicating phages with high numbers of tRNAs may be slightly more adapted to the host tRNA pool than phages with no tRNAs. The Low group had a significantly larger mean ΔtAI (−0.155) than the None group (−0.082, p-value 0.0257). In addition, the lifestyle of a phage did not seem to significantly affect how adapted a phage is to the host tRNA pool in *Bacillus* phages, though virulent phages show slightly lower levels of adaptation.

For the *Lactobacillus* phages, neither the logit-scaled intercept nor the comparison of lifestyle were significant. This indicates that temperate phages and those with no tRNAs do not significantly differ in tRNA adaptation from the host; thus the type of infection cycle does not significantly affect how well-adapted a phage is to its host tRNA pool in *Lactobacillus* phages either. However, both tRNA group comparisons showed significant differences between the High vs. None groups (mean ΔtAI of -0.147 and -0.036 respectively, p-value 0.0201) and the Low vs. None groups (mean ΔtAI of -0.129 and -0.036 respectively, p-value 0.0486). For these phages, having tRNAs in general is associated with lower levels of adaptation.

The *Staphylococcus* phages had a significant logit-scaled intercept (−0.333, p-value 3.01 x 10^-09^), indicating the temperate and no tRNA phages have significantly lower adaptation to the tRNA pool compared to the bacteria. Unlike the other Gram-positive hosts, the lifestyle comparison showed highly significant differences in mean ΔtAI for the temperate (−0.047) and virulent (−0.141) phages (p-value 3.23 x 10^-12^), indicating virulent *Staphylococcus* phages are significantly less adapted to the host tRNA pool than temperate phages. The tRNA group comparison for the *Staphylococcus* phages could only be performed with the Low vs. None groups, as none of the phages infecting *Staphylococcus* in this study have more than 4 tRNAs. These data indicate that the Low group (mean ΔtAI -0.141) is significantly less adapted to the host tRNA pool than the phages with no tRNAs (mean ΔtAI -0.047, p-value 3.23 x 10^-12^). The lifestyle and tRNA group comparisons were perfectly confounded, with all temperate phages having 0 tRNAs and all virulent phages belonging to the Low group.

In *Mycobacterium* phages, the only significant result was found for the intercept, with a logit-scaled value of -0.810 (p-value 3.98 x 10^-43^). These results suggest that *Mycobacterium* phages in general show significantly less adaptation to the host tRNA pool than the host. Neither the lifestyle comparison nor the tRNA group comparison yielded significant results, indicating that the lower adaptation of the *Mycobacterium* phages is a general feature, rather than specific lifestyle or tRNA characteristics of the phages.

### ΔtAI Comparison Globally Across Genera

Like the ΔtAI comparisons within a host genus, the global comparison across genera used a mixed model to assess the impact of phage lifestyle and number of tRNAs on how much a phage’s tAI differs from its host’s. This assesses the global trends in phage adaptation to their host tRNA pools based on the predictors of tRNA group and lifestyle. As before, the intercept was calculated using temperate phages and phages with no tRNAs as the baseline ΔtAI values. The logit-scaled intercept was -0.372, indicating significantly lower adaptation for temperate phages and those with no tRNAs (p-value 1.39 x 10^-13^). When tRNA group comparisons were run, the High group had a mean ΔtAI that was significantly larger than the mean ΔtAI of the None group (−0.158 vs. -0.088, p-value 2.31 x 10^-08^), indicating that globally, phages with high numbers of tRNAs have lower adaptation to their host tRNA pools. Likewise, the Low group had a mean ΔtAI of -0.135, which was also significantly larger than that of the None group (p-value 0.0074). On a global scale, lifestyle also had a significant impact on the difference between phage and host adaptation to the host’s tRNA pool. Virulent phages had a mean ΔtAI of -0.148, significantly larger than that of the temperate phages, which had a mean ΔtAI of -0.106 (p-value 0.0315).

Finally, on a global scale, the number of tRNAs, having tRNAs in general, and the lifestyle of a phage measurably affect how well-adapted a phage’s codon usage is to its host tRNA pool. The distribution of ΔtAI values for phages globally for both tRNA group comparisons and lifestyle comparisons are available in **Figure 13**. Specifically, phages encoding high numbers of tRNA genes, encoding tRNA genes in general, and being virulent are all associated with lower adaptation to the host tRNA pools. It is possible that phage-encoded tRNAs compensate for this by adding their own tRNAs to the host tRNA pool. These additions would change the composition of the tRNA pool inside the cell to one that is more favorable for their codon biases. Given that virulent phages have a finite amount of time to replicate and assemble new particles, this may provide a fitness benefit, considering that most of the phages carrying larger numbers of tRNAs are virulent.

**Figure 13:**
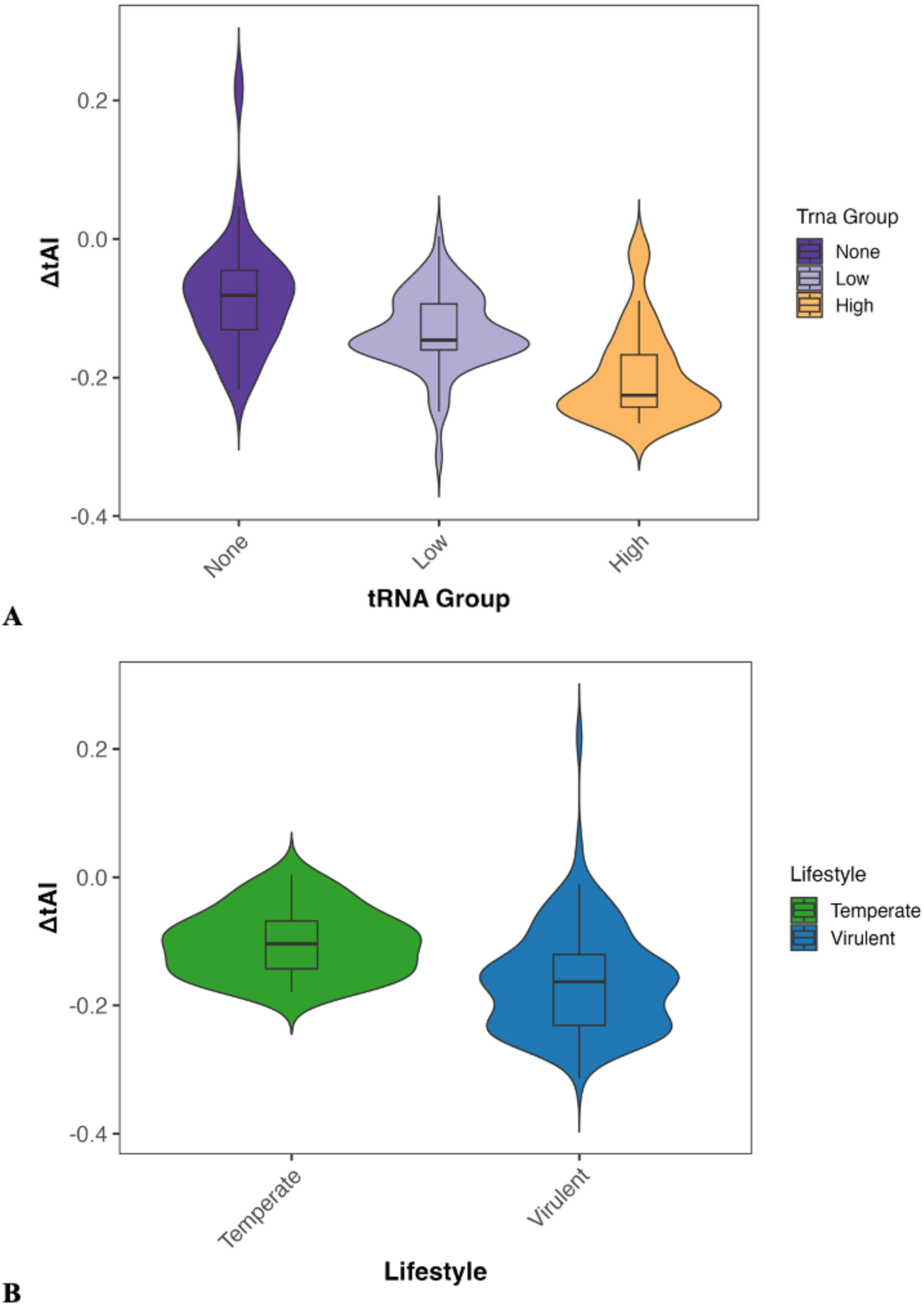
Distribution of global ΔtAI in phages. A) A violin plot depicting the global distributions of ΔtAI values in the phages belonging to each of the three tRNA groups (High, Low, None). High is colored in yellow, Low in light purple, and None in dark purple. B) A violin plot depicting the global distributions of ΔtAI values in the phages belonging to each lifestyle (Virulent or Temperate). Virulent are colored in blue, temperate in green.

## CONCLUSIONS

There is an incredibly complex relationship between the lifestyle, number of tRNAs, and codon usage bias of a phage and how it relates to that of its host. This work investigated that relationship, comparing codon usage between each phage and its respective host with multiple indices (GC content, EnC, RSCU, and tAI), then comparing phages by lifestyle and number of tRNAs both within host genus and globally across genera. The global trends indicate that virulence and high numbers of tRNA genes are associated with higher codon bias, with greater differences in codon bias and genomic composition compared to that of their hosts. At the genus level, these results are not consistent: phages infecting Gram-negative bacteria tended to have patterns more similar to the global trends compared to phages infecting Gram-positive bacteria. In phages infecting Gram-positive bacteria, there is less consistency across phages infecting different host genera, and in general these phages encode fewer tRNA genes compared to phages infecting Gram-negative hosts. Additionally, *Mycobacterium* phages tend to have different patterns altogether when compared to phages infecting other Gram-positive hosts, those infecting Gram-negative hosts, and global trends. So far, many of the large-scale studies investigating tRNA genes and codon usage have been performed in *Mycobacterium* phages. This work addresses these differences to define patterns for other bacteria and phage groups, providing information on a variety of host genera in addition to global trends.

## ACKNOWLEDGEMENTS

This work was supported by the National Institude of Allergy and Infectious Disease of the National Institutes of Health under award R01AI170608. The content is solely the responsibility of the authors and does not necessarily represent the official views of the National Institutes of Health.

